# Genetics, energetics and allostery during a billion years of hydrophobic protein core evolution

**DOI:** 10.1101/2024.05.11.593672

**Authors:** Albert Escobedo, Gesa Voigt, Andre J Faure, Ben Lehner

## Abstract

Protein folding is driven by the burial of hydrophobic amino acids in a tightly-packed core that excludes water. The genetics, biophysics and evolution of hydrophobic cores are not well understood, in part because of a lack of systematic experimental data on sequence combinations that do - and do not - constitute stable and functional cores. Here we randomize protein hydrophobic cores and evaluate their stability and function at scale. The data show that vast numbers of amino acid combinations can constitute stable protein cores but that these alternative cores frequently disrupt protein function because of allosteric effects. These strong allosteric effects are not due to complicated, highly epistatic fitness landscapes but rather, to the pervasive nature of allostery, with many individually small energy changes combining to disrupt function. Indeed both protein stability and ligand binding can be accurately predicted over very large evolutionary distances using additive energy models with a small contribution from pairwise energetic couplings. As a result, energy models trained on one protein can accurately predict core stability across hundreds of millions of years of protein evolution, with only rare energetic couplings that we experimentally identify limiting the transplantation of cores between highly diverged proteins. Our results reveal the simple energetic architecture of protein hydrophobic cores and suggest that allostery is a major constraint on sequence evolution.

## Introduction

A defining feature of globular proteins is the presence of a well-ordered hydrophobic core ^1–4^. Indeed the exclusion of water by the burial of hydrophobic side chains - the hydrophobic effect - is considered the major driving force in protein folding ^5–10^, and buried core residues are both highly conserved during evolution and very sensitive to mutation ^11–14^. In contrast, solvent-exposed residues on the surfaces of proteins are faster evolving with mutations typically having much smaller effects on stability ^1,15–20^. Surface residues can, however, be important for function, for example forming binding interfaces.

Many fundamental questions about protein hydrophobic cores remain unanswered. In particular, even for the smallest globular domains there are vast numbers of sequence combinations possible in the core-forming positions and the complexity of the mapping between sequence (genotype) and protein stability and function (phenotype) is poorly understood. At one extreme it could be that the dense packing of side chains in cores results in very complicated genotype-phenotype maps, with many pairwise and higher order combinatorial dependencies between the amino acids permitted at the different sites ^21,22^ making evolution hard to predict and engineering challenging ^23–29^. At the other extreme, mutations in cores might have largely independent effects, for example if core side-chain packing is highly malleable, allowing re-packing in many different combinations ^30–39^. Unfortunately, quantifying the effects of individual mutations ^15–17,19,20,40–42^ or pairs of mutations ^18,43–45^ provides little information about the genetic and energetic architecture of cores as it only explores very local sequence space, revealing the outcome when one or two side chains are changed. Rather, what is needed are experiments where the side chains of many buried core positions are simultaneously changed in many different combinations, an approach referred to as combinatorial mutagenesis or core randomisation ^3,46–53^. However, to date, the number of core genotypes that have been experimentally characterized is extremely small for any protein ^38,54–57^, meaning the complexity of protein core genotype-phenotype maps remains unclear.

A second fundamental question about hydrophobic cores is the degree to which the sequence of a core affects protein functions beyond stability. Although side chains buried internally in proteins might be assumed to have little effect on functions mediated via their surfaces, pioneering studies have identified cases where core mutations disrupt surface functions ^18,45,58–60^. These long-range ‘allosteric’ effects can be mediated by both propagated conformational changes and entropic effects ^61–63^.

Here, we use an experimental design in which we randomize the hydrophobic cores of proteins to better understand their genetic architecture, energetics, allostery and evolution. The resulting datasets reveal that the genetic and energetic architectures of protein cores are remarkably simple. This simplicity means that hydrophobic cores can be successfully transplanted between proteins separated by hundreds of millions of years of evolution, with only rare energetic couplings affecting stability. However, our data also reveal that many internal mutations in proteins have energetic consequences that propagate to the surface. The energetics of these allosteric effects is also simple, but their pervasive nature predicts that allostery has a large impact on sequence evolution.

## Results

### Randomizing the hydrophobic cores of proteins

To better understand the genetics, energetics and evolution of protein cores, we randomized the buried core residues of three structurally diverse proteins. For each protein we constructed a library in which seven core residues were randomized to five hydrophobic amino acids: F, I, L, M and V (Fig. 1b). Each library therefore contains 78,125 (=5^7) sequences, each with a different combination of hydrophobic residues in the core-forming positions.

**Fig. 1.**
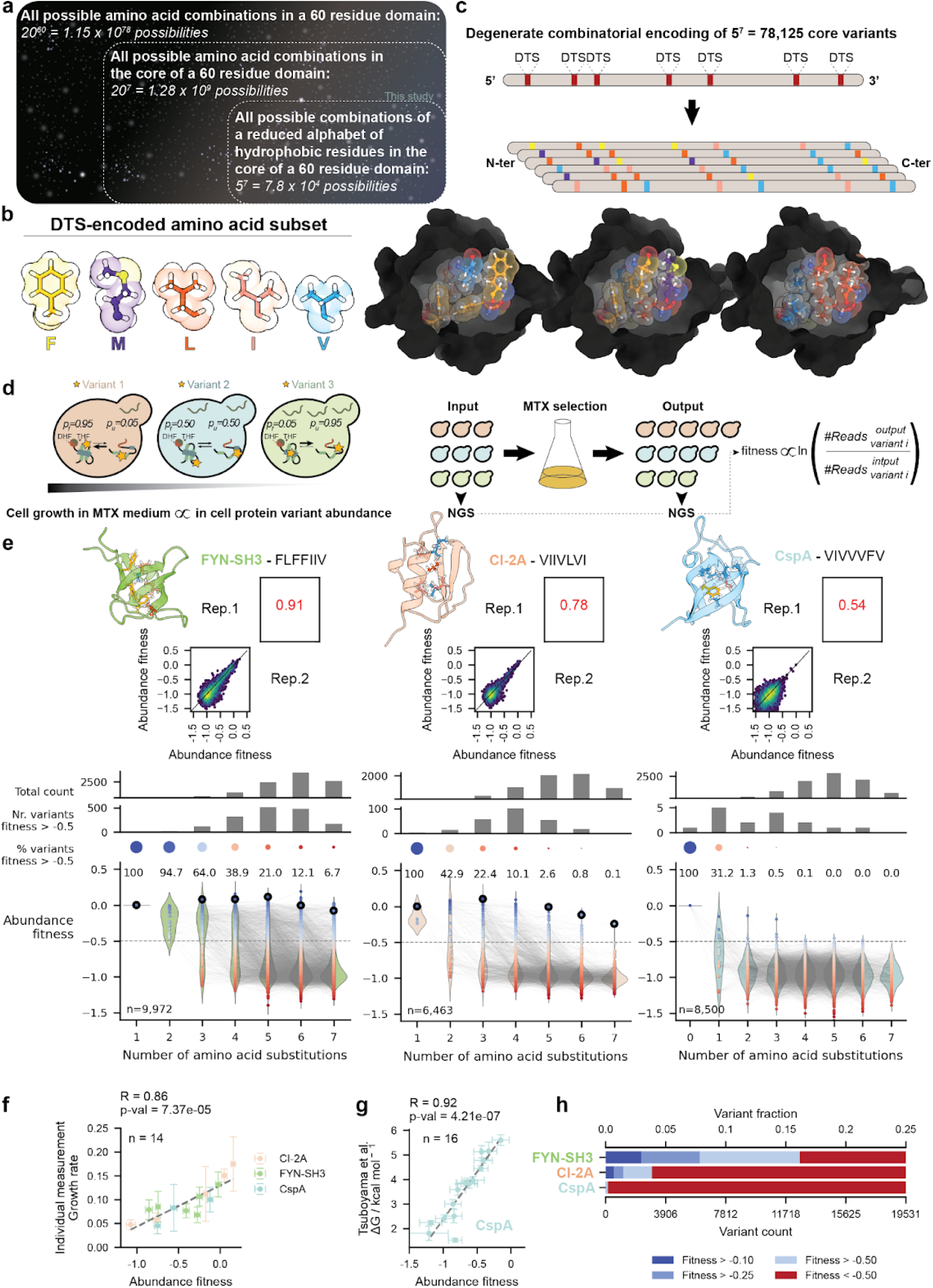
Randomizing protein hydrophobic cores at scale. **a.** Cartoon representation of the sequence space of an archetypal small protein with 7 core residues (arbitrary/not to scale). **b.** The subset of five hydrophobic amino acids encoded by the DTS degenerate codon. **c.** Hydrophobic core randomization is achieved by introducing the DTS codon in 7 core positions, yielding a 78,125 variant sequence space. **d.** In abundancePCA, the folding equilibrium of a given variant dictates the growth rate under selection of the cell population carrying that variant. Fitness measurements are proportional to the log ratio of NGS counts before and after selection. **e**. Abundance fitness measurement correlations in biological replicates and abundance fitness landscapes for each protein with a randomized hydrophobic core. Lines connect variants separated by 1 mutation. Circled variants were selected for *in vitro* characterization. **f.** Relationship between abundance fitness scores (± duplicate sigma) and individual clone growth rate measurements (± triplicate standard deviation). R, pearson correlation score. **g.** Relationship between abundance fitness scores (± duplicate sigma) and *in vitro* ΔG measurements (± CI95) ^19^. **h**. Fraction and count (extrapolated to the complete genotype space) of variants with abundance fitness above the indicated thresholds for each of the three proteins with randomized hydrophobic cores.

For each protein we measured the stability of a median of 8,500 randomly sampled genotypes using a highly-validated selection assay that quantifies the cellular concentration of folded protein over at least three orders of magnitude ^18,20,45,64,65^. The cellular protein abundance measurements were well correlated for each protein between replicate experiments (median Pearson correlation coefficient r=0.78, Fig. 1e). Abundance measurements also correlated very well with repeated measurements made for individual genotypes (r=0.86, n=14, Fig. 1f) and with independent *in vitro* measurements of fold stability (r=0.92, n=16, Fig. 1g).

### Stable cores

Considering all three proteins, 93.3% of core randomisations reduce abundance to less than 50% of the reference protein (wildtype or fittest lowest order variant; >50% abundance henceforth referred to as ‘stable’). However this varies across proteins, with 83.8, 96.1 and 99.8% of core randomisations having less than half the reference abundance for FYN-SH3, CI-2A and CspA, respectively (Fig. 1h). Despite the overwhelming detrimental nature of core mutations, in the full sequence space this means that >12,000 randomized FYN-SH3 cores, >3,000 CI-2A cores and >140 different CspA cores are stable. Consistent with expectations, proteins with only a single mutation are most likely to be stable, followed by those with two, three and then progressively more substitutions (Fig. 1e).

However, a large number of proteins with many substitutions in their cores still have high abundance. For example, for the FYN-SH3 domain, 94.7, 64.0, 38.9, 21.0, 12.1 and 6.7% of genotypes with one to seven substitutions in their cores have >50% reference variant abundance. Moreover, because there are many more genotypes with progressively larger numbers of substitutions (due to combinatorial explosion), most of the FYN-SH3 domains with high abundance actually contain many substitutions (Fig. 1e). Considering all 1,613 measured genotypes with >50% reference variant abundance, only one (0.1%) contains one substitution, 18 (1.1%) contain two, 114 (7.1%) contain three, 316 (19.6%) contain four, 513 (31.8%) contain five, 477 (29.6%) contain six and 174 (10.8%) contain seven substitutions. The solution space for stable hydrophobic cores can therefore be large and degenerate, with many solutions highly diverged from the wildtype protein.

### Combinatorial core mutants are folded and structured

We randomly picked examples of stable core mutants for two proteins, FYN-SH3 (Fig. 2a) and CI-2A (Fig. 2b), for further *in vitro* biophysical characterisation. We first performed thermal denaturation experiments. The wild-type proteins and all tested combinatorial mutants containing between one and all seven core residues mutated have sigmoidal denaturation curves, consistent with two-state cooperative folding. Moreover, the circular dichroism spectra of all combinatorial mutants match those of the wild-type proteins, consistent with conserved secondary structure content.

**Fig 2.**
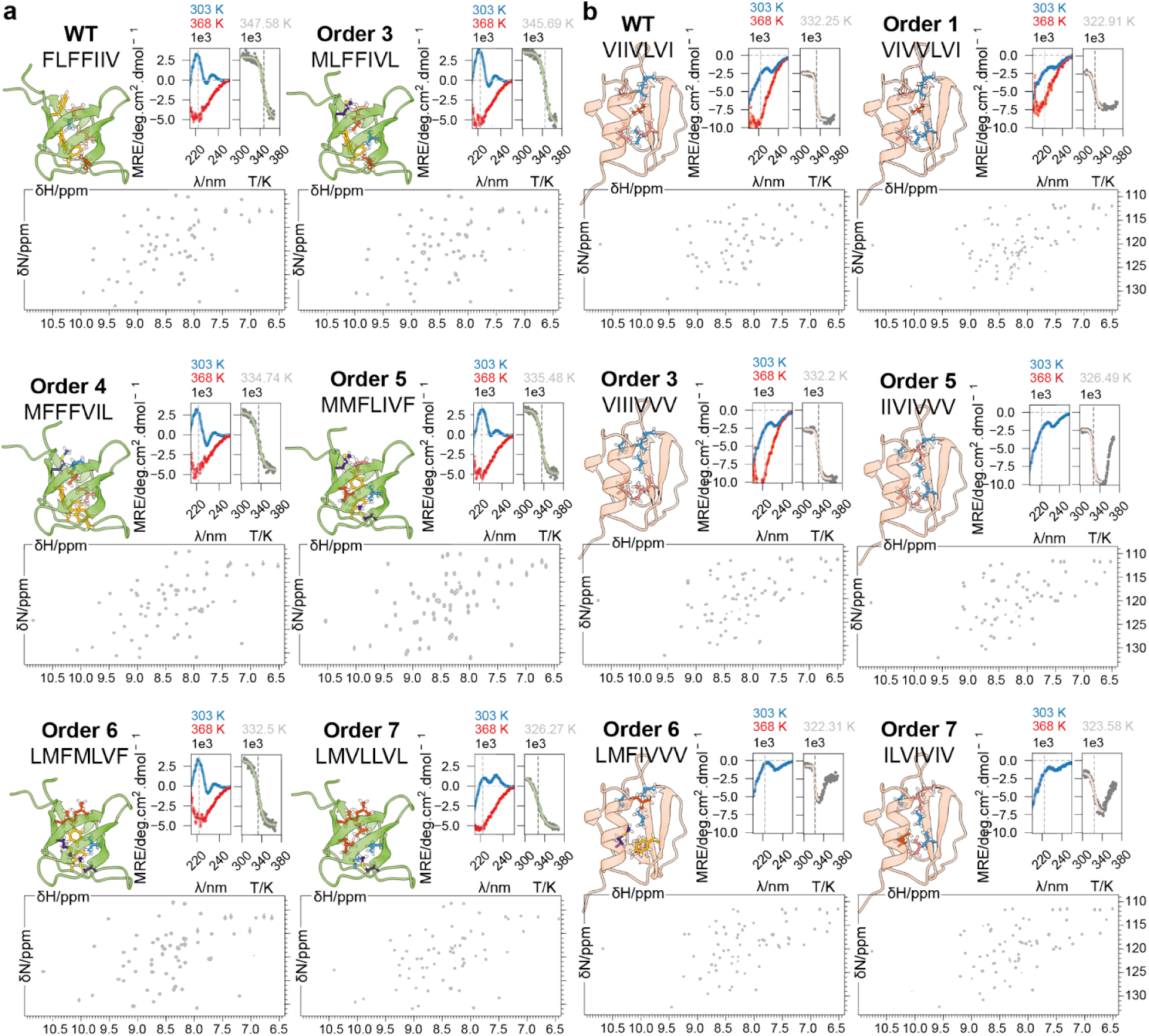
*in vitro* biophysical characterization of high order core variants. **a.** CD and NMR spectra acquired on recombinant samples of FYN SH3 WT and combinatorial core variants. AlphaFold2 models depict predicted core packings for each variant. Far-UV CD spectra are shown at the selection temperature (303 K, blue) and after denaturation (363 K, red), along with thermal denaturation curves acquired by monitoring MRE at 222 nm (T_m_ indicated with a gray dashed line). NMR ^1^H-^15^N HSQC spectra acquired at 303 K are shown for each variant. **b.** Same as in a. for CI-2A. CD spectra at 363 K are omitted for samples that collapsed and precipitated out of solution upon denaturation.

We next acquired ^1^H-^15^N heteronuclear single-quantum correlation (HSQC) nuclear magnetic resonance (NMR) spectra for the mutants and the wild-type proteins. All spectra showed the wide resonance dispersion in both dimensions and the sharp, well-resolved peaks expected of fully folded proteins. However, there are large ^1^H-^15^N HSQC resonance reconfigurations in the higher-order combinatorial core mutants, consistent with AlphaFold2 predictions of alternate core side chain packing (Figs. S1 and S2).

We conclude that, *in vitro*, these combinatorial mutants are folded and stable, but with altered side chain packing.

### Additive energy models predict core stability

The large solution spaces for stable hydrophobic cores present a challenge for predicting fold stability from sequence. A large solution space could arise from a complex genotype-phenotype map (‘fitness landscape’) with many pairwise and higher order epistatic interactions that would be hard to model, understand and represent in a compressed format ^23–29^. Such a landscape might arise, for example, in complex energetic interactions driving alternative side chain packing arrangements. Alternatively, the genotype-phenotype map could be quite simple, with each amino acid contributing an additive energy to fold stability.

We first tested a simple model in which mutations have phenotypic effects that combine additively. A linear regression model with four parameters per core position (one for each alternative aa) captures about 63.2% of the explainable variance in abundance across all random core proteins for FYN-SH3 and 42.9% for CI-2A (Fig. 3b). This result is striking, revealing simplicity in the genotype-phenotype landscape. The predictions of the additive phenotype models are, however, systematically biased (Fig. 3b, S3a). We therefore next tested a model in which mutations combine additively at the energy level, with changes in energy determining protein abundance by a sigmoidal function described by the Boltzman partition function for a two state cooperatively folding protein (Fig. 3a).

**Fig 3.**
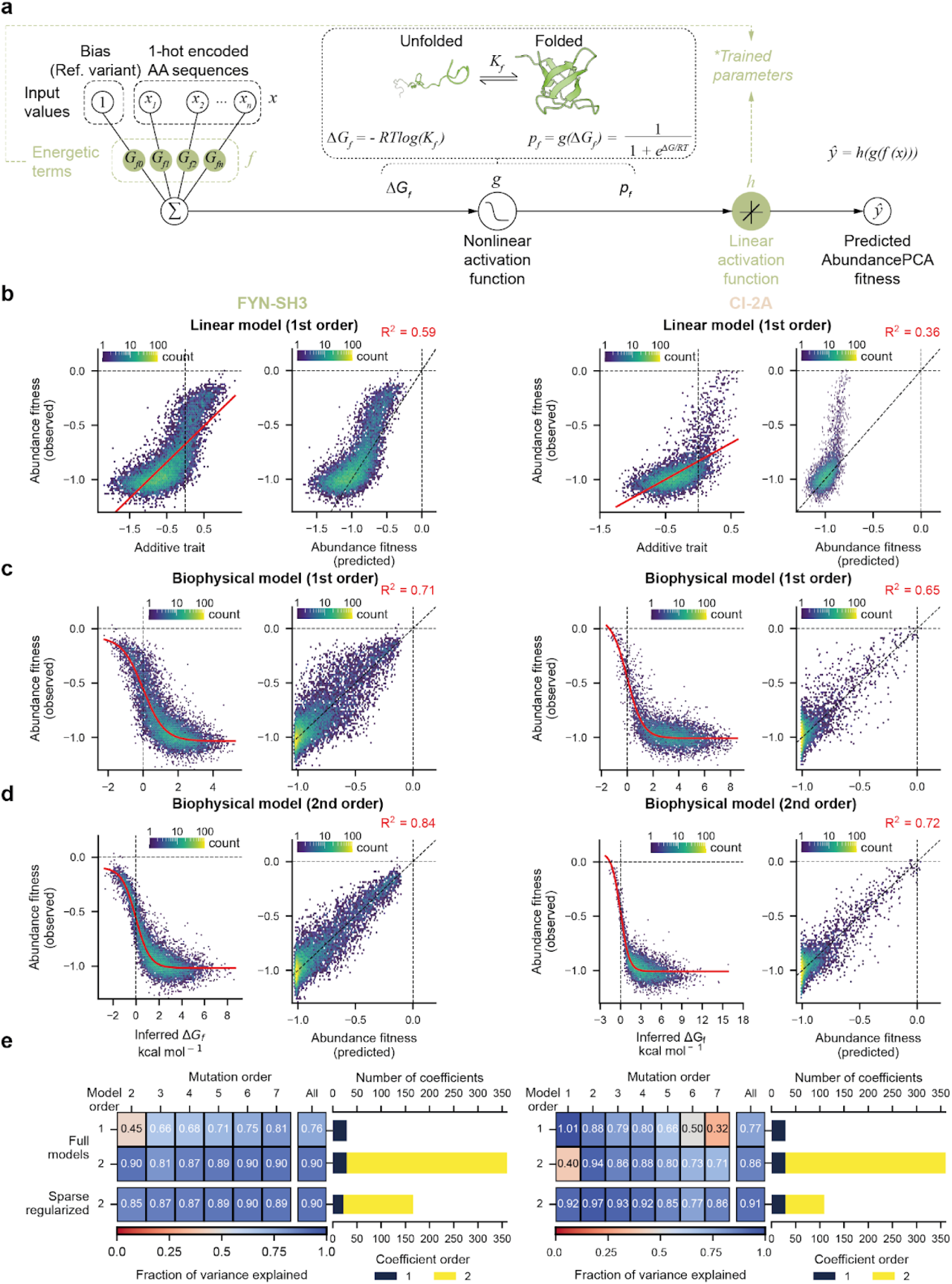
Energy models predict the stability of randomized protein cores. **a.** Architecture of the MoCHI neural network used to fit thermodynamic models (g) to the data (*ŷ*). *K_f_*, folding equilibrium constant; *ΔG_f_*, Gibbs free energy difference between the folded and unfolded states; *T*, assay temperature; *R*, gas constant; *p_f_*, probability of the folded state; *G_f_*, additive energy terms; *x_n_*, one-hot encoded variants; *h*, affine (shift and scale) transformation. **b.** Linear models assuming additivity at the phenotypic/ fitness level (solid red line) fit to the abundancePCA data (left panels), and corresponding model genetic prediction performance (right panels). **c.** First order two-state models assuming additivity at the energy level with no epistatic interactions. **d.** Second order two-state models accounting for pairwise energy couplings. **e.** Genetic prediction performance at all hamming distances evaluated by the fraction of variance explained (FVE) for all models fit to the data.

We used MoCHI ^66^ to fit energy models with a single additive change in Gibbs free energy of folding (ΔΔG_f_) for each mutation and found that these indeed provide improved predictive performance for FYN-SH3 (R^2^ = 0.71, fraction of variance explained (FVE) = 0.76) and CI-2A (R^2^ = 0.65, FVE = 0.77 for CI-2A; ten-fold cross validation). The additive energy models also largely eliminate bias (Fig. 3c, S3a). They contain 31 parameters (four energy terms per position, one for each mutation, totalling 28, one energy term for the wild-type and 2 global parameters relating the abundance fitness to the fraction of folded protein). They are therefore very large compressions (>2500:1) of the phenotype space.

### Sparse pairwise energetic couplings in protein cores

We next tested whether allowing mutations to energetically interact improved the performance of the energy models. With four mutations in seven positions there are a total of 336 (=7×4×6×4/2) pairwise energetic couplings (ΔΔΔG_f_) that can be quantified. Our experimental design provides very good power to quantify these couplings, for example each double mutant is tested in a median of 357 different genetic backgrounds in the FYN-SH3 dataset. Using MoCHI we found that models containing both the first order mutational effects (ΔΔG_f_, Fig 4a, b) and pairwise energetic couplings (ΔΔΔG_f_, Fig 4c, d) provided improved predictive performance for both FYN-SH3 (R^2^ = 0.84) and CI-2A (R^2^ = 0.72), approaching the upper limit of explainable variance estimated from the replicate correlations for these datasets (FVE = 0.90 and 0.86, respectively, Fig 3d, e). Moreover, many of the measured energetic couplings are very small (Fig. S4) and imposing sparsity using regularization eliminates many of the coupling energies without any loss in model performance (R^2^ = 0.84, FVE = 0.90 with 144 couplings and 169 total terms for FYN-SH3; R^2^ = 0.76, FVE = 0.91 for CI-2A with 80 couplings and 112 total terms for CI-2A, Fig. 3e, S3b). These second-order energy models are still large compressions of the phenotype space: >450-fold for FYN-SH3 and nearly 700-fold for CI-2A, and yet are sufficient for accurate reconstruction of the entire 78,125 genotype fitness landscape (Fig. 4e). We tested predictions of the model on unseen genotypes, further validating predictive performance (r = 0.85 for n = 77 variants measured in a new selection (FYN suppression library, see below), Fig 4g; and r = 0.73 for n = 10 individual clones, Fig. 4f).

**Fig 4.**
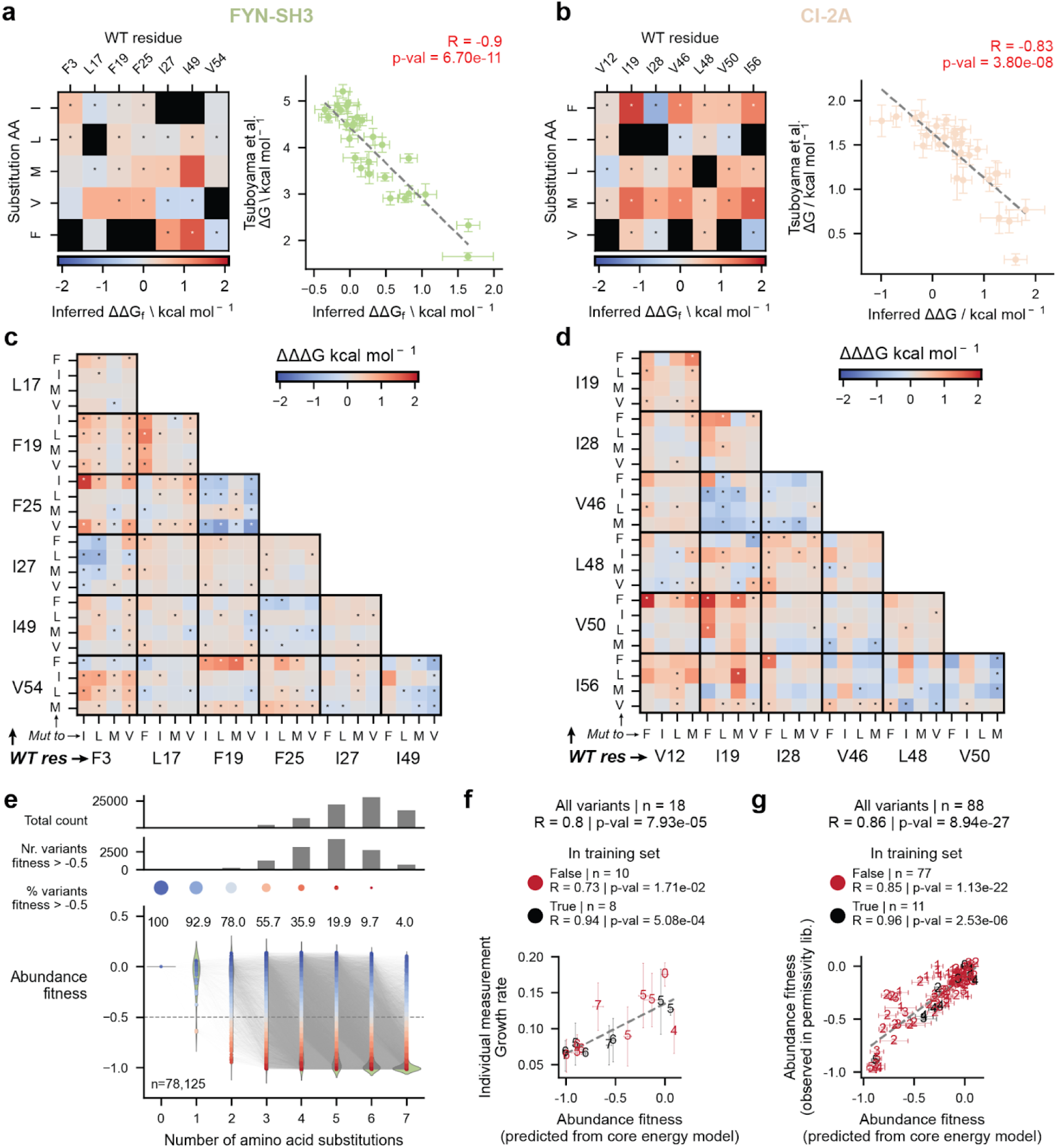
The energetic landscapes of two protein cores. **a.** Genetic background-averaged single mutant effects (*ΔΔG_f_*) inferred from the FYN-SH3 dataset by the full second-order biophysical model (left). Coefficients selected by the sparse regularized model are labeled with an asterisk (Fig. S4a,b). Right: relationship between inferred *ΔΔG_f_* (± std from 10-fold model cross-validation) and *in vitro ΔG_f_* values (± CI95) ^19^. *R*: pearson correlation score. **b**. As in **a** for the CI-2A core DTS dataset. **c.** Pairwise energetic couplings (*ΔΔΔG_f_*) in the core of the FYN SH3 domain inferred by the full second-order biophysical MoCHI model, with coefficients selected by the second-order sparse regularized model labeled with asterisks (Fig. S4c,d). **d.** Same as in **c** for CI-2A. **e.** Combinatorially complete abundance fitness landscape of the FYN SH3 DTS core predicted by the second-order energy model. **f.** Relationship between abundance fitness predictions (± 10-fold std) and individual growth rate measurements (± triplicates std) for variants present and not present in the model training set. R: pearson correlation score. **g.** Relationship of abundance fitness values predicted by the second-order energy model fit on FYN-SH3 core mutagenesis data (± 10-fold std) and abundance fitness values experimentally measured in the FYN-SH3 suppression library selection (± sigma, see below).

### Predicting core stability across hundreds of millions of years of evolution

Our randomization experiments quantified the ability of combinations of hydrophobic residues to constitute a stable buried core in one particular sequence context. For these particular contexts we were able to train highly predictive protein core energy models. Given the sparsity and simplicity of these models, we tested how well they perform in different sequence contexts beyond the proteins for which they were trained by assessing how well they predict the stability of core amino acid combinations naturally found in structural homologs (Fig. 5a). The Earth Biogenome provides one large-scale test of model performance across diverse sequence contexts. Using structural homology ^67^ we identified 51,159 unique SH3 domains across 2,545 species (Fig. 5b, S5). 15,304 of these SH3 domains have one of the hydrophobic residues F, I, L, M and V at all seven of the residues that we randomized in our FYN-SH3 library, identifying a combination of hydrophobic residues that is functional in the context of that SH3 domain sequence. The natural domains vary in their sequence identity to the FYN-SH3 domain (henceforth referred to as FYN) allowing us to quantify how well the FYN hydrophobic core energy model performs in predicting functional cores in SH3 domains with between 98.2 and 8.6% sequence identity.

**Fig 5.**
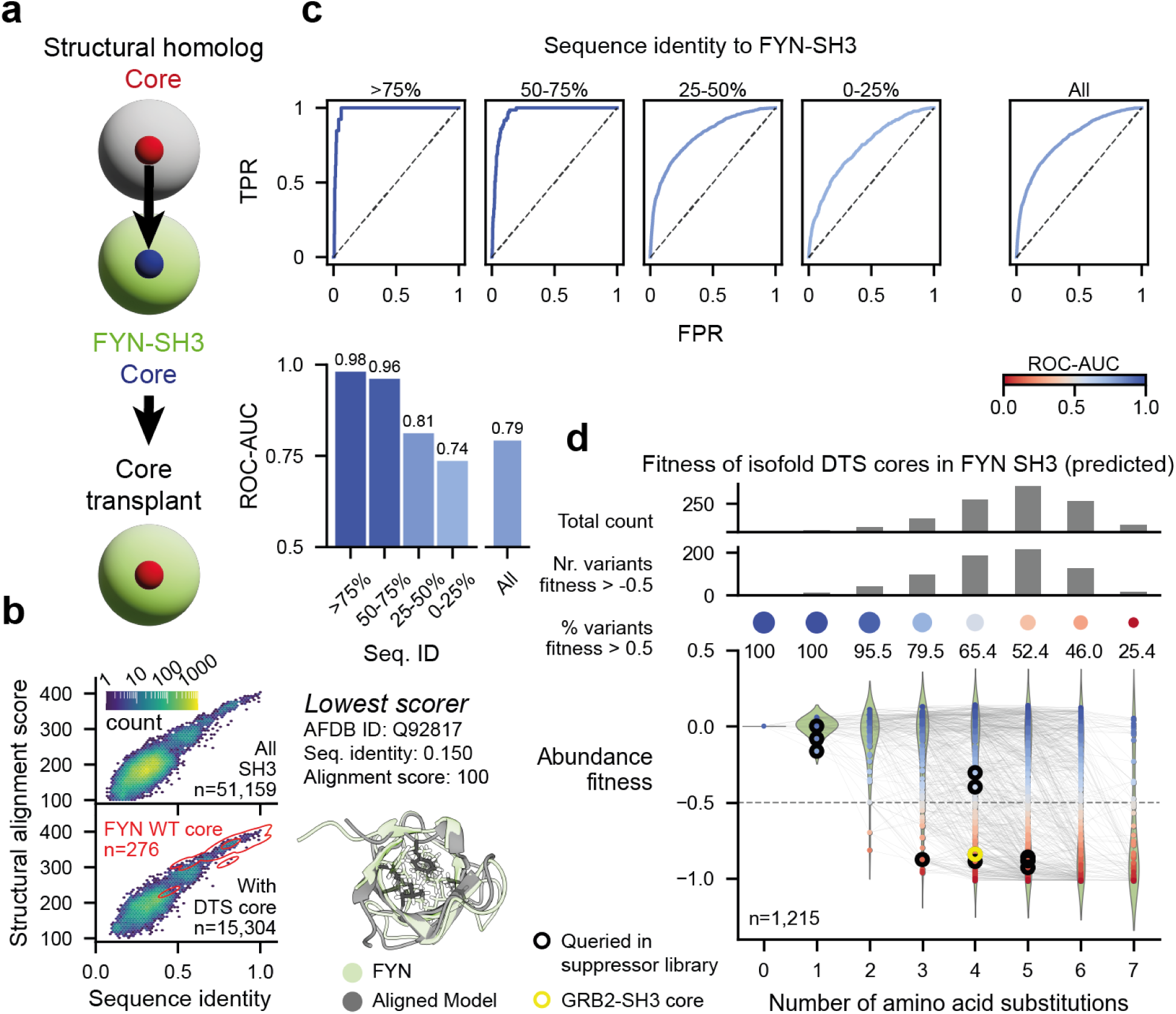
Predicting SH3 domain hydrophobic core stability across a billion years of evolution. **a.** Scheme of the core transplant concept to assess the performance of the energy models trained on FYN-SH3 core mutagenesis at predicting hydrophobic core stability across evolution. **b.** SH3 domains identified by structural homology as classified by their sequence and structural similarity to FYN-SH3. **c.** Receiver operating characteristic (ROC) curves for a classification task aimed at distinguishing between cores existing or not in the subsets of the identified structural homologs (top) based on the complete landscape (78,125 variants) of folding free energies predicted by the second-order energy model. ROC-AUCs are shown as gradient colored bars (bottom). **d.** Mapping in the FYN-SH3 core predicted abundance fitness complete landscape of the 1,215 unique F, L, I, M, V core combinations identified in the structural homologs set. Circled variants were selected for transplant failure recovery experiments (GRB2-SH3 core circled in yellow).

Strikingly, we find that the FYN core energy model is highly predictive across the entire SH3 domain family, with a receiver operating characteristic curve area under the curve (ROC-AUC) for classifying observed from non-observed hydrophobic cores in the test set equal to 0.79 for all 15,304 SH3 domains (Fig. 5c, S6a). Performance is extremely good for SH3 domains with high sequence similarity to FYN (ROC-AUC=0.98 for domains with >75% identity) but also very good for much more evolutionarily diverged SH3 domains. For example, performance is still excellent (ROC-AUC=0.96) for domains with between 50% and 75% identity to FYN. For more diverged domains, performance is reduced but still highly predictive with a ROC-AUC=0.81 for domains with 25-50% identity and ROC-AUC=0.74 for domains with very low (<25%) identity (Fig. 5c).

Thus, a model trained on data from the experimental mutagenesis of only one protein is able to predict functional hydrophobic cores across the entire sequence diversity of SH3 domains in the Earth Biogenome, an evolutionary timescale of at least one billion years ^68^.

### Sequence determinants of core transplantation failures

The reduced performance of the FYN energy model in the most distantly related SH3 domains suggests that combinations of hydrophobic residues that confer stability in FYN and closely related SH3 domains sometimes do not confer stability in more highly diverged SH3 domains.

To test this idea more directly, we used our data to measure whether core sequences observed in natural SH3 domains also confer stability when ‘transplanted’ into FYN (Fig. 5a). As shown in Fig. 5d, the FYN hydrophobic core energy model covers transplants into FYN of 1,215 distinct hydrophobic cores from 15,304 natural SH3 domains. In total, 57.4% of these transplanted cores confer stability sufficient to obtain >50% of the abundance of wild-type FYN. The remainder we refer to as ‘transplantation failures’.

To identify the causes of core transplantation failures (Fig. 6a), we focussed on eight combinations of hydrophobic amino acids that are frequently observed in natural SH3 domain cores (occuring in between 140 and 3,512 examples) but are detrimental when transplanted into FYN (each transplant destabilizes the protein by between 0.90 and 2.73 kcal mol^-1^, Fig. 6b). We confirmed the stability of natural structural homologs carrying these cores by quantifying the abundance of 62 WTs and 3 proline core mutants for each of them (Fig. 6c and S6b). To identify candidate sequence changes in other residues that allow these cores to confer stability in natural SH3 domains but not in FYN (Fig. 6a), we trained logistic regression models to distinguish SH3 domains with these cores from the remainder i.e. those without them (Fig. 6d). We then designed a FYN variant library to test whether each of the single AA substitutions highlighted by the regression models could – alone and in combination – rescue the detrimental effects of the core transplantation into FYN. In total, we tested between 17 and 21 single AA substitutions and between 234 and 363 combinations of variants for their ability to rescue each transplantation (Fig. S7). We also used a similar approach to test candidate suppressor mutations for 14 variants that are mildly detrimental in FYN but observed at high frequency in natural SH3 domains (Fig. S8).

**Fig 6.**
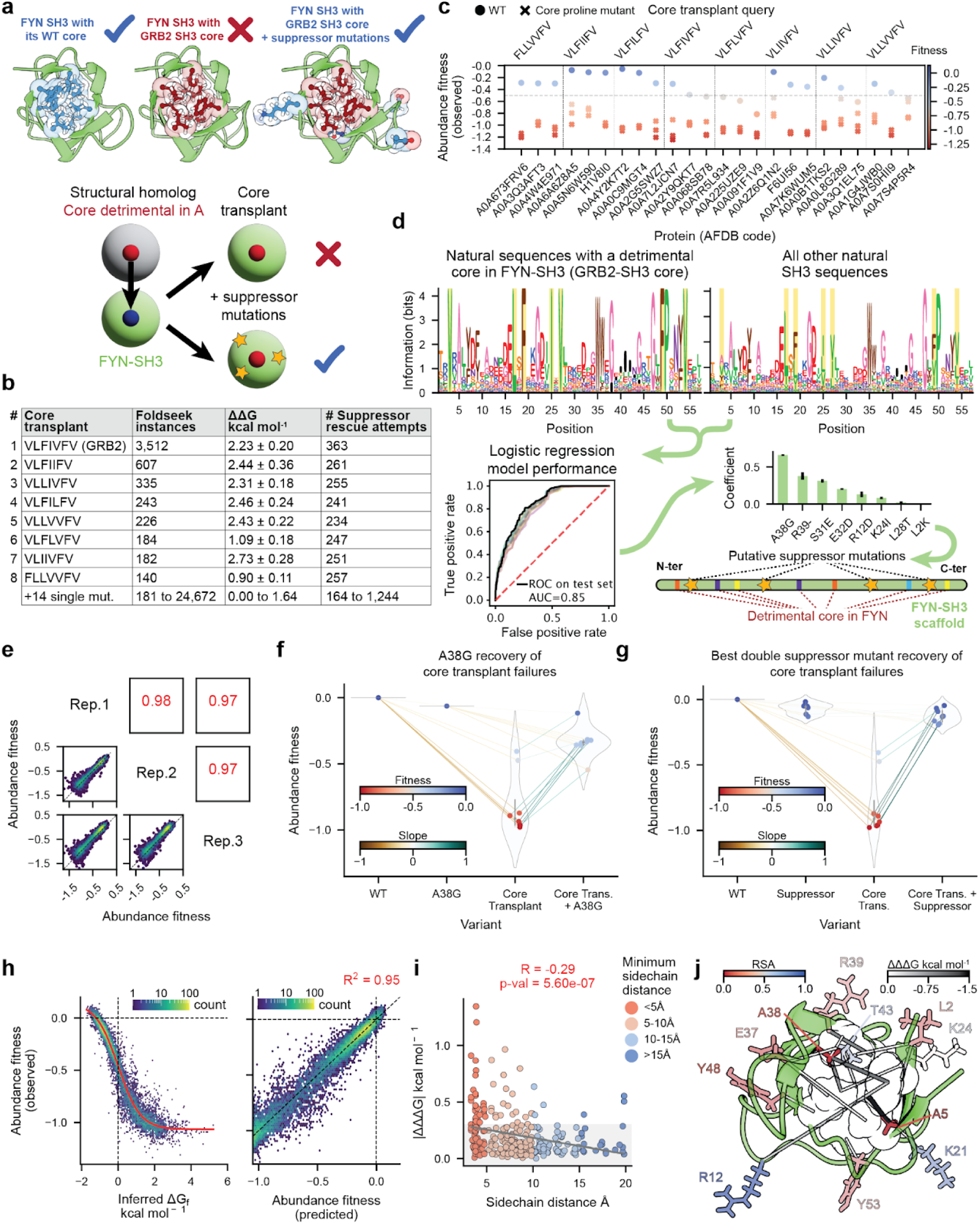
Determinants of core transplantation failures. **a.** Scheme of the suppression hypothesis for recovery of core transplant failures. **b.** Core transplant failures selected for recovery with suppressor mutations. **c.** Experimental measurements of abundance fitness of structural homologs carrying cores whose transplant in FYN fails (top 3 per core transplant query, full set in Fig S6b). **d.** Strategy to identify putative suppressor mutations. The subset of sequences presenting a particular query core combination (for example, the GRB2-SH3 core combination, in yellow shades) is compared to the rest of the sequences using a logistic regression model with one-hot encoded amino acid sequence features using the FYN WT sequence as the reference. Candidate suppressor mutations are identified as selected or highly ranked model coefficients in classifiers trained with and without regularization. **e.** Biological replicate correlations of the abundance fitness measurements on the FYN suppression library selection. **f.** A38G abundance fitness rescue and suppression of detrimental effects of failed natural core transplants in FYN. The observed fitness of all eight failed core transplants is shown, and each is connected to the fitness of the variant carrying both the core transplant and the A38G suppressor mutation. **g.** Best double suppressor mutation fitness recovery of core transplant failures. **h.** Regularized second-order model fit on the FYN suppression library selection data (left, red line) and associated genetic prediction performance (right). **i.** Relationship between pairwise coupling magnitudes and side chain heavy atom distances. Shaded area denotes the mean magnitude of couplings at <5Å. **j.** Stabilizing (<0 kcal mol^-1^) pairwise energy couplings involving a core residue (shown as transparent spheres) and a suppression site (shown in sticks) are depicted as solid lines on the structure of FYN SH3 (PDB 5ZAU). Side chains of coupled residues are colored according to their calculated RSA. Thickness and color of the lines represent the magnitude of the coupling.

This variant library contained 11,647 FYN variants, and abundance fitness measurements were highly reproducible (r ≥ 0.97, Fig. 6e). For all eight core transplants the library identified a mutation that, when introduced into FYN, strongly suppressed the detrimental effects of transplantation (Fig. S7). For 7/8 detrimental transplants the same mutation - A38G, a substitution in a buried residue (RSA = 0) - was the strongest individual substitution suppressor (Fig. 6f). For the other transplantation the strongest suppressor was D15N, a mutation in a distal (9.04 Å minimal side chain heavy atom distance to the core) partially buried position (RSA = 0.59) in loop 1. Suppression by these individual substitutions was strong but not complete. However, for all eight core transplantations a combination of two mutations (all including A38G) was sufficient to restore FYN stability to wild-type levels (Fig. 6g). Suppressors were also identified for all individual mildly detrimental core substitutions (Fig. S8).

We used MoCHI to fit a regularized energy model to the complete transplant-rescue dataset. The model was sparse (>19-fold compression, 604 terms for 11,647 genotypes) but highly predictive (R^2^ = 0.95 by ten-fold cross-validation, Fig. 6h), including 24 energies (ΔΔG_f_) for the mutations in the core and 146 for the mutations outside of the core (Fig. S9) and 431 pairwise energetic couplings ΔΔΔG_f_ quantified in a median of 102 genetic backgrounds (32 between core residues (7.4%), 256 between a core and a non-core residue (59.4%), and 143 between non-core residues (33.2%), Fig. S10) whose magnitude is proportional to the 3D distance between the coupled residues (Fig. 6i). The strongest suppressive energetic couplings (ΔΔΔG_f_ < 0 kcal mol^-1^) with mutations outside of the core are with mutations in sites packed against the core (A5, A38, Fig. 6j). However the model also includes couplings with partially buried sites in the core vicinity (L2, E37, R39, Y48, Y53) and some couplings with surface sites (R12, K21, T43), suggesting the potential for long-range indirect energetic couplings in this domain.

In summary, these test cases where the FYN-SH3 core-only energy model fails to accurately predict stability during long-term evolution can thus be explained by a small number of energetic couplings to mutations elsewhere in the protein domain, predominantly - but not exclusively - in residues proximal to the core.

### Stable cores frequently disrupt function

We next designed an experiment to test whether hydrophobic core randomization affects protein function beyond changes in stability. SH3 domains are protein interaction domains that recognise proline-rich sequence motifs through a flat binding interface on their surface ^69,70^. None of the seven core residues randomized in our experiments form part of this binding interface (Fig 7a,b) ^71–73^. To test whether FYN-SH3 domains with randomized hydrophobic cores are functional we quantified their binding to an engineered high-affinity 14-residue peptide variant from the Csk-binding protein (Cbp), a transmembrane adaptor protein activated in T-cells by FYN phosphorylation upon MHC engagement ^73,74^. We quantified the binding of 45,480 randomized core variants to the peptide ligand (Fig. 7c-e). Ligand binding measurements were highly reproducible (median r = 0.92 across three replicate experiments, Fig. 7d).

**Fig 7.**
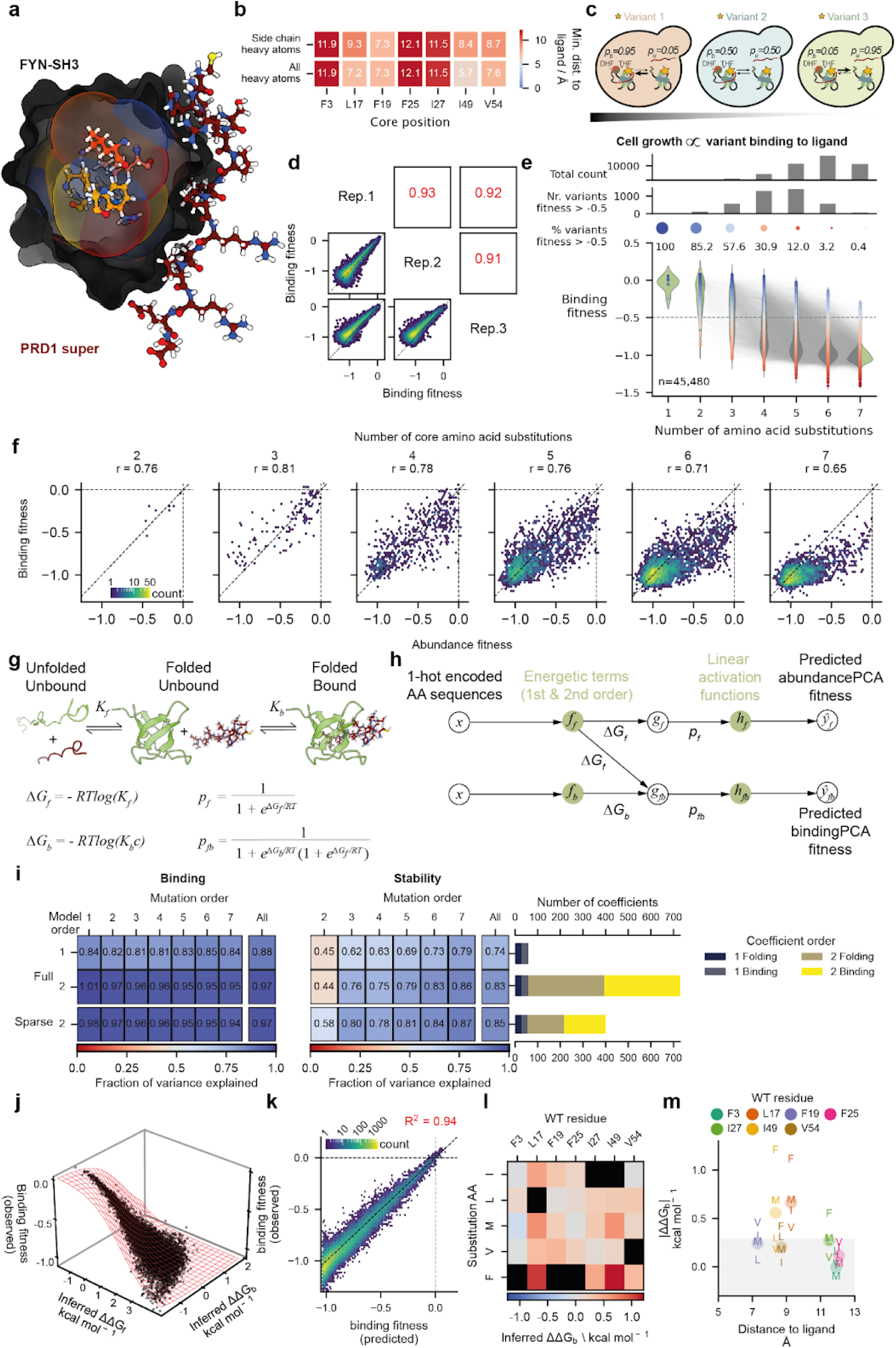
Pervasive allostery constrains hydrophobic core sequence evolution. **a.** AlphaFold2 model of the interaction between the FYN-SH3 domain and the PRD1 super peptide. **b.** Heavy atom minimum distances between mutagenized core positions and the peptide ligand. **c.** Scheme of the bindingPCA readout used to obtain binding fitness measurements. **d.** Biological replicate correlations of binding fitness measurements on the FYN randomized core library binding to PRD1 super. **e.** Binding fitness landscape, with variants separated by mutation order and connected by a line if one mutation apart. **f.** Relationship between abundance and binding fitness measurements for each mutational order. r: pearson correlation score. **g.** Three-state thermodynamic model of the folding and binding equilibria used to simultaneously fit abundance and binding fitness data. **h.** Architecture of the neural network used to fit the three-state model to the data. **i.** Genetic prediction performance at all hamming distances evaluated by the fraction of variance explained (FVE) for all models fit to the data. **j.** Relationship between the observed binding fitness values and the inferred free energies of folding and binding (data as black dots, sigmoidal fit shown as a red mesh). **k.** Genetic prediction performance of the second-order three-state model on binding fitness. **l.** Genetic background-averaged binding free energy changes (ΔΔG_b_) associated to FYN-SH3 core point mutations inferred by the second-order three-state model. **m.** Magnitude of individual (characters) and position averaged (dots) single mutational binding free energy effects (|ΔΔG_b_|) versus minimum side chain heavy atom distance to ligand.

In total, 3,896 FYN-SH3 variants (>6,600 when extrapolated to the complete mutational landscape) with randomized hydrophobic cores bound the ligand (binding defined as >50% of reference FYN-SH3 high fitness variant). This provides additional strong evidence that many FYN-SH3 domains with randomized cores are correctly folded.

However, plotting the relationship between ligand binding and abundance for proteins with different numbers of mutations in their cores revealed a surprising relationship: as the number of core mutations increases, the effects on binding appear greater than the effects on folding (Fig. 7f), suggesting that mutations in the core might be having indirect i.e. allosteric effects on the binding affinity.

### Extensive allostery in hydrophobic protein cores

To better quantify the allosteric effects of sequence changes in the core of FYN-SH3 we used MoCHI to fit a three state energy model to the combined binding and abundance datasets (Fig. 7g,h). In this model, mutational effects on both the folding free energy (ΔG_f_) and ligand binding free energy (ΔG_b_) are inferred, with free energy changes combining additively in combinatorial mutations (Fig. 7i and Fig. S11). This simple additive energy model provides very good predictive performance for both binding (R^2^ = 0.85, FVE = 0.88) and abundance (R^2^ = 0.69, FVE = 0.74; ten-fold cross validation). The model is a nearly 900-fold compression of the data (62 parameters to generate 55,452 phenotypes). The additive energy model correctly predicts that higher order combinatorial core mutations frequently have larger effects on ligand binding than protein abundance (Fig. S11i), suggesting that the impaired binding in combinatorial core mutants is mainly driven by additive energy changes. Although individually quite mild, the pervasive allosteric effects of individual mutations (Fig. S11d) add together in combinatorial mutants to strongly impair binding. One consequence of this is the larger effects of core transplantation experiments on binding than on folding (Fig S12).

### Accurate prediction of the allosteric effects of core randomisation

To investigate a role for energetic couplings within the core in allostery, we fitted a more complex energy model that quantifies the pairwise energetic couplings for folding (ΔΔΔG_f_) and binding (ΔΔΔG_b_) (Fig. 7i-l). This model improves prediction of the binding (R^2^ = 0.94, FEV = 0.97, Fig. 7i,k) and abundance (R^2^ = 0.77, FEV = 0.83, Fig S13c) data. Imposing sparsity through regularization reduces the complexity without affecting performance (R^2^ = 0.94, FEV = 0.97 in binding and R^2^ = 0.79, FEV = 0.85 in folding; 404 parameters: >130-fold compression, Fig. 7i and S14), suggesting that many of the energetic couplings are effectively zero.

The improved predictive performance when including energetic couplings suggests an important - albeit limited - role for epistasis in the effects of core randomisation on FYN-SH3 binding. Just as for stability changes, most changes in binding upon the randomisation of protein cores can be accounted for by additive changes in free energy. It is the sum of many individually mild and moderate allosteric effects (Fig. 7m) that can prove catastrophic in combination.

## Discussion

We have quantified here the stability and ligand binding of tens of thousands of proteins with randomized hydrophobic cores. Our experimental design deliberately limited randomisation to hydrophobic amino acids, allowing us to generate a very large dataset of stable cores with highly diverged sequences. Fitting energy models to this data revealed that the fundamental genetic and energetic architecture of protein cores is strikingly simple: both protein stability and ligand binding can be accurately predicted over very large evolutionary distances using additive energy models with a limited contribution from pairwise energetic couplings. These energy models are simple and overlook fine-details of folding and binding equilibria ^75^ yet they capture most of the variance in the data. This simplicity extends to the interaction of core residues with the rest of the protein, allowing cores to be successfully transplanted between proteins separated by hundreds of millions of years of evolution, with only rare - and identifiable - energetic couplings reducing stability.

We have also shown here that many combinations of internal mutations in proteins have large energetic consequences that propagate to the surface. The allosteric effects of combinatorial mutants arise not because of complicated epistatic fitness landscapes but rather because of the pervasive nature of allostery: many individual core mutations have mild or moderate allosteric effects and these sum together to produce consequential binding energy changes in combinatorial mutants. Such pervasive weak allostery will have important evolutionary consequences, predicting that sequence changes in cores may frequently be selected against because of their effects on function rather than stability ^60^.

The simplicity of hydrophobic core fitness landscapes and pervasive allostery have important implications for protein engineering. Our results - and the data of others ^39,76,77^ - show that protein stability can be predicted and engineered using simple and interpretable models. This should accelerate protein engineering and design and make it more understandable and explainable. Understanding and engineering allosteric effects is a major challenge and directed evolution campaigns often select for unexpected allosteric mutations outside of active sites ^78,79^. Our results show that allosteric mutations can be frequent and that their combined effects can be quite straightforward to predict. Additional experiments with diverse protein folds and functions will be required to evaluate the generality of these conclusions.

Perhaps most strikingly, our results show that vast numbers of amino acid combinations can constitute stable protein cores but that these alternative cores frequently disrupt protein function because of allosteric effects. However, our results also show that the genetics and energetics of both core stability and allostery are quite simple and interpretable, allowing accurate prediction over very large evolutionary timescales and adding to a growing body of recent work ^39,76,77^ suggesting that proteins are rather less mysterious and complex than is widely perceived.

## Data availability

All DNA sequencing data have been deposited in the Gene Expression Omnibus under the accession number GSE266299. Files to reproduce all figures in this work are available at: https://doi.org/10.5281/zenodo.11175470

## Code availability

Source code used to perform all analyses and to reproduce all figures in this work is available at: https://github.com/lehner-lab/combinatorialcores

## Author contributions

A.E. and GV performed experiments. A.E., G.V and A.J.F. performed analyses. A.E. and B.L. conceived the project, designed the experiments and analyses, and wrote the manuscript, with input from A.J.F.

## Acknowledgements

This study was funded by a European Research Council (ERC) Advanced (883742) grant, the Spanish Ministry of Science and Innovation (LCF/PR/HR21/52410004, EMBL Partnership, Severo Ochoa Centre of Excellence), the Bettencourt Schueller Foundation, the AXA Research Fund, Agència de Gestió d’Ajuts Universitaris i de Recerca (AGAUR, 2017 SGR 1322), the CERCA Program/Generalitat de Catalunya and Wellcome (Grant reference: 220540/Z/20/A, ‘Wellcome Sanger Institute Quinquennial Review 2021-2026’). We thank all members of the Lehner Lab at the CRG and the Wellcome Sanger Institute for their valuable feedback and the Laboratory of Molecular Biophysics led by Prof. Xavier Salvatella at the IRB Barcelona, the Barcelona node of the R-LRB NMR facility and the CCTiUB at the University of Barcelona for assistance with NMR and CD spectroscopy data acquisition.

## Materials and methods

### Core mutagenesis library design

The DTS codon was identified by iteratively searching degenerate codons encoding subsets of hydrophobic amino acids with minimal degeneracy using the python package dogma (https://pypi.org/project/dogma/). Residues to mutagenize in FYN-SH3, CI-2A and CspA were prioritized by K-means clustering all protein residues based on their relative accessible surface area (RSA) along with their closeness centrality in the structure contact network. The latter was built on the basis of contact maps calculated on experimental PDB structures, with a heavy-atom distance cut-off of 5Å to assign a contact between a residue pair. Clustering was performed in each model of the NMR structures with PDB accession numbers 5ZAU (FYN-SH3), 3CI2 (CI-2A), 2L15 and 3MEF (CspA). 7 residues per protein were then prioritized based on the fraction of models where they were clustered as core residues, their belonging to the F, L, I, M, V set and their tight packing in the structure. In FYN-SH3, positions 3, 17, 19, 25, 27, 49 and 54 were selected; in CI-2A, positions 12, 19, 28, 46, 48, 50 and 56; and in CspA, positions 5, 17, 26, 28, 47, 49 and 63.

### Core stability in structural homologs across the Earth Biogenome and FYN suppression library design for core transplant failure recovery

To identify FYN-SH3 structural homologs across the Earth Biogenome, we used Foldseek ^67^ to align the 5ZAU PDB structure against the AlphaFold database ^80^. Foldseek returned 435,428 results that were filtered to ensure core positions mutagenized in FYN were structurally aligned and thus comparable to the Foldseek-aligned positions in the structural homologs. Filters required that the homologs be aligned at least to positions 2 to 54 in FYN-SH3 (covering all core positions), have an alignment TM-score > 0.6, a C_α_ RMSD < 2 Å, and an alignment probability of query and target being homologous (same SCOPe superfamily) = 1. 131,696 structural homologs found in 2,622 different species passed the filters, constituting 51,159 unique amino acid sequences belonging to > 2,500 species. Of these, 43,111 structural homologs had cores with all seven positions being F, L, I, M or V (15,304 unique sequences after removing duplicates). 276 structural homologs with no duplicated sequences had the same core configuration found in FYN-SH3.

The performance of the energy models trained on the sparsely sampled FYN-SH3 DTS core library at predicting the existence of the entire genotype space (5^7 = 78,125 possible core amino acid combinations) was evaluated as follows. Genotypes were binarily-labeled depending on whether they were found or not in the Foldseek-identified set of structural homologs (or a subset defined by sequence identity to FYN-SH3) which constituted the target variable. The classification performance of the folding free energy (ΔG_f_) predicted for each genotype by the energy model was evaluated by the receiver operating characteristic area under the curve (ROC-AUC).

Natural core combinations with detrimental effects in FYN-SH3 stability were selected as queries in the FYN suppression library based on the number of sequences with a particular combination of amino acids in their cores (n = 140 to 3,512) and on the destabilizing severity of the high order mutations when transplanted to FYN-SH3 as predicted by the 2nd order energy model fit on the core mutagenesis abundancePCA data (ΔG_f_ ranging from 0.90 ± 0.11 to 2.73 ± 0.28 kcal mol^-1^). 14 core point mutants were also selected for suppression mutation rescue based on their background-averaged destabilizing effect inferred from the same model.

Candidate suppressor mutations to attempt fitness rescue of the core transplant failures and detrimental point mutations were identified via sequence comparison by training a logistic regression model. An MSA was built from Foldseek pairwise structural alignments against FYN-SH3, whose sequence was used as the reference and no insertions were allowed to avoid misalignments. One-hot encoded sequences were used as features for the regression task.

Query core positions were completely masked out (all seven positions originally mutagenised in FYN-SH3 for natural core query transplants, just the one mutant position for core single mutant queries). Uninformative, low-variance features with positional frequencies ≤5% and ≥95% were also masked out. The presence/absence of the query core configuration was used as the target variable. Sequences were randomly assigned to training (90%) and test (10%) sets, and the regressor was trained with 10-fold cross-validation both with and without Lasso regularization. In the feature selection approach with regularization, we iteratively decreased the L1 penalty with the aim of retaining at least 6 non-zero features (up to 9 depending on the query). We then encoded suppressor mutations as all combinatorial possibilities in the resulting binary space. In the feature selection approach without regularization, we selected the top 15 model coefficients ranked by magnitude and encoded suppressor mutations as all possible single and double mutant combinations in the binary space. The final library totalling 12,000 variants also included 10 natural isofold sequences for each of the 8 natural core queries, that is, 80 sequences corresponding to natural proteins carrying the cores that are detrimental when transplanted to FYN-SH3, and three random proline core mutants for each of them.

### Library construction and cloning

Oligonucleotides, plasmids and media used are listed in Tables S1,S2 and S3, respectively. Combinatorial core libraries using a reduced alphabet of hydrophobic amino acids (F, L, I, M, V) were built by multiple segment Gibson assembly of oligonucleotides bearing the DTS degenerate codon at core residue-encoding positions (oligos used in this study are detailed in Table S1 and were all obtained from IDT, Coralville, IA, US). Each of the genes encoding the FYN SH3 domain, CI-2A and CspA were codon optimized for expression in *Saccharomyces cerevisiae* and divided in two segments smaller than 200 bp each, which was the size limit for commercial synthetic degenerate oligo production at the time of performing this study. Both the 5’ and the 3’ gene segments contained a homology region of at least 25 bp to the linearized pGJJ191 mutagenesis vector, plus an internal homology region of at least 25 bp. Double stranded Gibson assembly inserts were produced from single stranded commercial oligonucleotides via a single cycle PCR reaction using a 3’ annealing reverse primer specific to each segment (see Table S1). The pGJJ191 vector was linearized via a 25-cycle PCR reaction using primers oGJJ311 and oGJJ406, followed by 1h DpnI (NEB) treatment at 37°C plus inactivation at 80°C for 20 minutes (1 μL enzyme per 50 μL PCR reaction). Both inserts and vector PCR reactions were cleaned-up in MinElute columns (Qiagen, Hilden, Germany). Gibson assembly mixtures contained 300 ng of linearized pGJJ191 plus the two insert segments at a 5:1 molar ratio each, along with a Gibson enzyme mixture produced in house by the Protein Technologies core facility at the CRG.

Conversely, the FYN SH3 suppression library was purchased as an oligo pool from Twist Bioscience (San Francisco, CA, US). The oligo pool was amplified in a 13 cycle PCR reaction with primers oAE018 and oAE227, which was cleaned up using a MinElute column. The insert amplified oligo pool contained 25 bp homology regions at both ends and was mixed at a 5 to 1 molar ratio with 300 ng of linearized pGJJ191 and subsequently with the Gibson enzyme mixture.

All Gibson assembly reactions were incubated at 50°C for 8h, dialyzed against water using 0.025 μm filters (Millipore, Burlington, MA, US) for 1.5h, concentrated using a SpeedVac and transformed into *E. coli* C3020 cells (NEB, Ipswich, MA, US). The number of transformants was assessed by colony count in serial dilutions of transformation outgrowth aliquots in LB plates supplemented with 50 μg/mL spectinomycin, and 1 to 5 equivalent transformation reactions were pooled to reach at least 20X variant coverage per library as required. Plasmid DNA was extracted using the Qiagen Plasmid *Plus* Midi kit and the library inserts were obtained through double endonuclease digestion (HindIII-HF and NheI-HF, NEB) followed by agarose gel purification achieved by a combined use of centrifugal filter units (Millipore) and MinElute clean-up. Inserts were introduced in double-restricted, Quick CIP-trated (NEB) pGJJ162 (abundancePCA) or pGJJ159-PRD1super (bindingPCA) vectors by overnight thermal-cyclic ligation using T4 DNA ligase. The pGJJ159-PRD1super vector was previously obtained by introducing the *S. cerevisiae* codon-optimized PRD1super encoding gene from commercial oligo synthesis (IDT) into BamHI-SpeI double restriction-linearized pGJJ159 via Gibson assembly. Ligation mixtures were dialyzed against water, SpeedVac-concentrated and transformed into *E. coli* C3020 electrocompetent cells. Equivalent transformations were combined to reach at least 20X variant coverage, and selection-ready libraries in PCA vectors were Midi-prep extracted.

For sparse sampling of combinatorial core libraries (FYN SH3, CI-2A and CspA in pGJJ162), the full-complexity libraries were transformed into *E. coli* C3020 electrocompetent cells and bottlenecked by outgrowth medium serial dilution into overnight selective medium (LB supplemented with 50 μg/mL ampicillin) assessed in parallel through colony count of plated aliquots, aiming at ∼10,000 transformants per library. Bottlenecked libraries were extracted by mini-prep, qPCR quantified and mixed to equivalent molar ratios.

### Large-scale transformation and competitive selection

The bottlenecked library pool containing randomly subsampled fractions of the FYN SH3, CI-2A and CspA DTS core libraries in pGJJ162, the full complexity FYN SH3 DTS core library in pGJJ159-PRD1super and the FYN SH3 suppression library in pGJJ162 were transformed in BY4741 yeast cells by heat shock. The number of transformation reactions was scaled depending on the final selection volume, which was in turn scaled to achieve a 100X coverage at the most stringent selection step in terms of cell count bottlenecking (inoculation in selection medium, see below). An individual transformation was performed per 200 mL of selection culture. For each individual transformation, 25 mL YPDA were inoculated with BY4741 colonies from a freshly streaked YPDA plate and grown overnight at 30°C. The process was performed in parallel for each independent biological replicate, starting from a different BY4741 colony (2 replicates for the subsample DTS core library pool, 3 replicates for the other experiments). The precultures were used to inoculate 180 mL of YPDA to an OD_600_ of 0.3, which was grown for 4 h at 30°C. Cells were harvested by 5 min centrifugation at 3,000 g, washed in water and treated with 8.6 mL SORB for 30 min. 3.5 μg of plasmid DNA were then added along with 175 μL of 10 mg/mL sonicated salmon sperm DNA (Santa Clara, CA, US), and the mixture was agitated at room temperature for 10 minutes followed by addition of 35 mL of plate mixture plus another 30 min incubation. 3.5 mL of DMSO were then added and the mixture was heat-shocked at 42°C for 20 min. After the heat-shock, cells were harvested by 5 min centrifugation at 3,000 g, resuspended in recovery media and incubated at 30°C for 1 h. Cells were then harvested again and resuspended in 200 mL of SC -URA selective medium. A 10 μL aliquot was serially diluted and plated in SC -URA to assess transformation efficiency.

All biological replicate transformation cultures were grown in parallel at 30°C for ∼24 h. Then, the selection input cultures were inoculated by resuspending water washed harvested cell pellets in SC -URA -ADE to an OD_600_ of 0.1 (800 mL in 5 L flasks in the cases of the subsampled DTS core library pool and FYN SH3 binding library, 400 mL in 2L flasks in the case of the FYN SH3 suppression library). The input cultures were grown at 30°C and harvested upon reaching an OD_600_ of 1.6 (5 generations), which typically occurred from 12 to 15 h after inoculation. Cell pellets were stored at −20°C. Aliquots of the input culture were used to inoculate the selection (output) cultures (SC -URA -ADE, 200 μg/mL methotrexate) at an OD_600_ of 0.05. The output cultures were grown at 30°C for ∼30 h (subsampled DTS core library pool and FYN SH3 binding library) or until they reached an OD_600_ of 1.6. At that time, cells were harvested by 5 min centrifugation at 3,000 g, thoroughly washed in water and stored at −20°C.

### DNA extraction, plasmid quantification and sequencing library preparation

Input and output culture cell pellets were resuspended in DNA extraction buffer (10 mM Tris-HCl, 100 mM NaCl, 2% Triton-X, 1% SDS, pH 8). Volumes were scaled depending on the volume of cell pellets to process. For pellets from 100 mL cultures harvested at OD_600_ 1.6, 1 mL of DNA extraction buffer was used. Cells were lysed by cyclically freezing in liquid nitrogen followed by incubation in a 62°C water bath twice. Total DNA was extracted by adding 1 mL of a 25:24:1 phenol:chloroform:isoamyl alcohol mixture (Merck, Darmstadt, Germany) along with 1 g of acid washed glass beads (Sigma-Aldrich, Saint Louis, MO, US) and vortexing for 10 min. The mixture was centrifuged at room temperature for 30 min at 3,300 g and the aqueous phase (upper layer) was recovered in a separate tube. 1 mL of the phenol:chloroform:isoamyl alcohol was added to it and the mixture was vortexed for 2 min followed by 45 min centrifugation at 3,300 g. The aqueous phase was again transferred to a new tube and mixed with 0.1 volumes of 3 M sodium acetate and 2.2 volumes of pure ethanol previously cooled down at −80°C. The mixture was kept at −20 °C for 30 min and spun down at 3,300 g for 30 min at 4°C. The DNA pellets were dried overnight in a fume extraction hood.

Plasmid DNA from total DNA extracts was quantified by qPCR using the oGJJ152-oGJJ153 oligo pair that anneals at the origin of replication of all pGJJ vector series. The qPCR reaction was performed in a LightCycler 400 instrument (Roche, Basel, Switzerland) using the SYBR green qPCR 2X master mix from Thermo Fisher (Waltham, MA, US). Quantification was achieved by measuring a standard curve on a serial dilution of a control sample of known concentration in the same qPCR run as the query samples, and subtracting blank measurements.

Library preparation for sequencing in Illumina instruments followed a two-step PCR protocol. Frameshifts and a segment of the Illumina adapter overhangs were introduced in PCR1. For each sample, 8×50 μL PCR1 reactions were performed using 31.25 million plasmid molecules as a template as quantified by qPCR. PCR1 used the oGJJ595 and the oGJJ748 frameshifting oligo pools (Table S1) and consisted of 11 cycles. Next, all reactions corresponding to the same sample were pooled and treated with 16 μL ExoSAP for 30 min at 37°C, then for 20 min at 80°C. Samples were cleaned-up using MinElute columns, eluted in 30 μL of pre-warmed water (55°C) and 1.5 μL were used as template for each PCR2 reaction.16×50 μL 8-cycle PCR2 reactions were performed per sample to introduce the rest of the Illumina adapters and the dual index barcodes. PCR2 reactions corresponding to the same sample were pooled, concentrated using a MinElute column, and run in a 2% agarose gel. Bands of the correct sizes were excised from the gel and sequencing-ready DNA was recovered by using a centrifugal filter unit followed by MinElute clean-up.

### Next-generation sequencing data acquisition, processing and analysis

Illumina NGS-ready samples were quality controlled using a TapeStation instrument (Agilent, Santa Clara, CA, US), quantified by qPCR and paired-end sequenced in either a NextSeq500 (FYN SH3 suppression library) or NextSeq2000 (subsampled core DTS library pool and FYN SH3 binding library) instrument (Illumina, San Diego, CA, US) by the CRG Genomics core facility. Raw reads were de-multiplexed by leveraging 5’ and 3’ unique dual indices introduced to each sample during PCR2.

Raw FastQ files were processed using DiMSum v1.3 ^81^ (https://github.com/lehner-lab/DiMSum) using default settings with minor adjustments. Experimental design files and command-line options required for running DiMSum on these datasets are available on GitHub (https://github.com/lehner-lab/combinatorialcores). Reads were retained if corresponding to variants expected by library design as specified in the *barcodeIdentityPath* argument and had a minimum Q score of 20 across all bases as specified by the *vsearchMinQual* argument. Variants were retained if accounted for by a threshold number of reads in all biological replicates chosen for each experiment to discard low-read variants and specified by the *fitnessMinInputCountAll* argument. Absolute growth rates were inferred by DiMSum from initial and final OD_600_ measurements and selection times for each replicate.

Fitness estimates for each variant *i* in replicate *r* were obtained as follows:

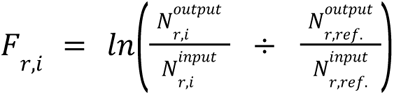

where 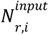 and 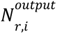 are the number of input and output reads for variant *i* in replicate *r* in the input and output samples, respectively; and *ref.* is a reference variant. Fitness measurement errors (*sigma)* were obtained by fitting the DiMSum error model to the data ^81^.

For data visualization and comparison, raw DiMSum fitness measurements were scaled to fall in the −1 to 0 range by centering the lower mode of the fitness values distribution to −1 and the fitness of the reference variant to 0. The reference variant for each dataset was defined as the lowest order, fittest variant that passed the filters mentioned above. Associated fitness errors (*sigma)* were also scaled by multiplying the raw values by the same scaling factor as the fitness values.

### Thermodynamic modeling

Linear and biophysical models were fitted to the raw fitness measurements and error estimates obtained from DiMSum using MoCHI ^66^. MoCHI is a neural network framework for fitting interpretable models to DMS data. In Fig. 3a, the global epistasis function (*g*) corresponds to a simple two-state folding model, where the equilibrium constant *K_f_* defines the difference in free energy between the folded and unfolded states (*ΔG_f_*) at a given temperature *T* provided the gas constant *R*. The probability of the folded state *p_f_*is related to *ΔG_f_* by the Boltzmann distribution. In this model, coefficients of the additive trait map (*f*) represent first-order (*ΔΔG_f_*) and pairwise (second-order) energetic coupling terms (*ΔΔΔG_f_*) applied to one-hot encoded sequence variants (*x*). An affine transformation *h* maps the molecular phenotype (fraction of molecules folded) to the measurement scale of the target variable (growth rate) ^66^.

AbundancePCA datasets for FYN-SH3 and CI-2A from the sparse core DTS library selection were fit with MoCHI either specifying no global epistasis (“*Linear*” model i.e. assuming mutant effects sum additivity at the phenotypic level) or specifying a two-state model of the folding equilibrium (“*TwoStateFractionFolded*” i.e. assuming additivity at the energy level). First- and second-order models were fit to the data (*max_interaction_order* command line argument), with the latter accounting for pairwise energetic couplings. Second-order models were simplified by imposing coefficient sparsity through regularization with search over 3 different L1 regularization penalty values (MoCHI command line arguments *l1_regularization_factor “0.01,0.001,0.0001”*). As stochastic gradient descent algorithms do not give exact zeros for coefficients with L1 regularization we used a greedy stepwise feature selection strategy that (i) starts by fitting a full model including all possible coefficients of all orders, (ii) refits the model without the highest order coefficients (i.e. setting them to zero) that are not nominally significant in step i, and (iii) repeats steps i and ii progressively for lower order sets of coefficients until (and including) coefficients of first order (MoCHI command line argument *sparse_method “sig_highestorder_step”*).

Similarly, a regularized second-order two-state model was fit to the abundancePCA data obtained from the FYN-SH3 suppression library using an equivalent configuration.

The bindingPCA data obtained for the FYN-SH3 DTS core variant library binding to the PRD1-super ligand was jointly fitted together with the corresponding FYN-SH3 abundancePCA data from the sparse core DTS library selection using a three-state model of the folding and binding equilibria (Fig. 7g). In this case, the neural network architecture consisted of two additive trait layers (Fig. 7h), one applying the *TwoStateFractionFolded* non-linear (global epistasis) transformation on the abundancePCA data relating each variant’s fraction of folded molecules with the underlying folding free energy, and the other one applying the *ThreeStateFractionBound* non-linear transformation on the bindingPCA data simultaneously relating each variant’s fraction of bound molecules with the underlying binding and folding free energies. Again, full first and second-order models and a regularized second-order model were fit to the data.

All MoCHI models were trained using 10-fold cross validation and model performance was evaluated on the unseen (held-out) data. We used MoCHI v0.9 (https://github.com/lehner-lab/MoCHI) to fit all models described here. Model design files and command-line options required for running MoCHI on these datasets are available on GitHub (https://github.com/lehner-lab/combinatorialcores).

The fraction of explainable variance (FEV) was obtained by dividing the maximum explainable variance (MEV) by the total fitness variance:

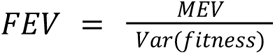

MEV is defined as the total fitness variance minus the total estimated technical variance as reported by DiMSum fitness errors:

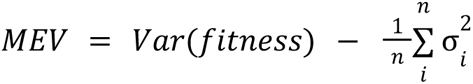

where σ_i_ is the error estimate sigma for variant *i* as obtained from DiMSum error model. The fraction of explained variance (FVE) was then obtained by dividing the R^2^ correlation score on observed *versus* predicted held-out data by FEV.

### Individual clones growth rate measurement

To isolate individual clones of FYN-SH3, CI-2A and CspA core variants present and covering the full dynamic range of fitness values in the sparse DTS core library pool selection, a 2 μL aliquot of one of its output DNA extracts was transformed into *E. coli* 10-beta cells (NEB). Instead, to obtain individual clones of FYN-SH3 core variants not measured in the experiment, the full non-bottlenecked library was transformed into *E. coli* 10-beta cells. Transformations were serially diluted and plated on LB plates supplemented with 50 μg/mL ampicillin. For screening, isolated individual colonies were inoculated in 96 well plates loaded with 400 μL LB + ampicillin per well. The mini-prep pre-cultures were sealed with a gas-permeable lid and grown overnight at 37°C. Cells were harvested by centrifugation (3,200 g, 10 min), resuspended in 25 μL P1 buffer (Qiagen) and lysed with P2 buffer. After 5 minutes, lysis buffer was neutralized with buffer P3, and cell debris was separated from the cleared lysate by centrifugation (3,200 g, 15 min). 60 μL of the supernatant were transferred to a new 96 well-plate loaded with 80 μL ice cold isopropanol per well. The plate was incubated at −20°C for 20 minutes and centrifuged again. After removing the isopropanol, 80 μL of 70% ethanol in water were added per well and DNA pellets were washed by pipetting. After ethanol removal, plates were dried overnight in a fume hood. Dry DNA was dissolved by adding 25 μL of water per well, and clones were screened by Sanger sequencing using oligo oGJJ083.

1.5 μL of selected individual clone mini-preps were transformed into 10 μL aliquots of BY4741 cells previously treated to become competent as detailed above (large-scale transformation and competitive selection methods section) and flash-frozen in liquid nitrogen after SORB treatment. The mini-prepped DNA was mixed with pre-boiled sonicated salmon sperm DNA (2 μL at 1 mg/mL) and 40 μL of plate mixture.The mixture was added to the cells, which were subsequently incubated at room temperature for 30 min, added 4 μL of DMSO and heat-shocked at 42°C for 20 minutes in a PCR block. Cells were harvested by centrifugation (3,000 g, 5 min) and resuspended in 100 μL recovery medium. After incubating at 30°C for 1h, 2.5 μL were spread on SC -URA plates and incubated at 30°C for 48h.

Individual yeast colonies were inoculated in a 384 deep well plate loaded with 150 μL of SC -URA -ADE per well, which was incubated overnight at 30°C with thorough agitation. After that, cells were resuspended by pipetting and diluted by transferring 5 μL of the saturated pre-cultures to a new Nunc 384 well flat-bottom plate (Thermo Fisher) loaded with 95 μL of water per well. Selection cultures were inoculated by pipetting 5 μL of the diluted cells into 75 μL of SC -URA -ADE disposed in a Nunc 384 well flat-bottom plate. Cell growth was monitored in a Tecan Infinite 200 Pro M Plex microplate reader with temperature control at 30°C by recording the OD_600_ every 15 min for 300 cycles (75 h).

### Recombinant isotope-labeled protein sample production

^15^N isotope-enriched FYN SH3 and CI-2A wild-types and selected variants were recombinantly produced and purified for *in vitro* characterization. Variants were selected at nearly all hamming distances based on their associated abundance fitness values, which were required to be in the higher end of the fitness values distributions at each hamming distance. Sequences were codon optimized for expression in *E. coli* and obtained in two separate synthetic oligo pools from IDT. Each pool was separately cloned into a linearized pDEST17-His-Sumo vector via Gibson assembly. Double stranded insert DNA was obtained from single-cycle PCR reactions on the synthetic oligo pools using oligo oAE208. The vector was linearized using primers oAE206 and oAE207. Gibson reactions were performed using 300 ng of the vector along with inserts at an insert:vector molar ratio of 5 to 1. The reactions were incubated at 50°C for 8 h and 1 μL of the mixture was then used for chemical transformation in NEB 10-beta competent cells. Transformation reactions were plated in LB plates supplemented with 50 μg/mL ampicillin and colonies were screened using Sanger sequencing.

Selected clones were transformed into BL21(DE3) chemically competent *E. coli* cells (NEB) and individual colonies were used to inoculate 50 mL overnight LB cultures grown at 37°C. Cells were harvested and used to inoculate 1L of M9 minimal medium with ^15^N ammonium chloride as the sole nitrogen source (Eurisotop, Saint-aubin-des-bois, France). Cultures were grown to an OD_600_ of 0.4 at 37°C with 220 rpm agitation, then transferred to a 25°C incubator and further grown to an OD_600_ of 0.6. At that stage, protein expression was induced by the addition of 0.5 mM isopropyl β-D-1-thiogalactopyranoside (IPTG) which proceeded overnight at 25°C with 220 rpm agitation.

Cells were harvested by centrifugation at 3,000 g for 15 min at 4°C and resuspended in cell lysis buffer (20 mM Tris-HCl, 150 mM NaCl, 20 mM imidazole, pH 8.0) supplemented with 1.5 mg/mL chicken egg white lysozyme, 0.1 mg/mL bovine pancreas DNAseI (both from Sigma-Aldrich), 0.5 mg/mL phenylmethylsulfonyl fluoride (PMSF) protease inhibitor and a tablet of EDTA-free Pierce protease inhibitors (both from Thermo Fisher). Cell lysis was performed by homogenization at a maximum 1,500 psi using an Emulsiflex-C5 instrument (Avestin, Ottawa, ON, Canada). Cell debris was removed by ultracentrifugation (40,000 g, 20 min, 4°C) and the cleared lysates were loaded on a 5 mL HisTrap Fast Flow column (Cytiva, Marlborough, MA, US) equilibrated in cell lysis buffer and mounted on an Äkta Pure system (also from Cytiva) placed in a 4°C refrigerator cabinet. After loading, the column was thoroughly washed in 5 column volumes of lysis buffer, 3 column volumes of wash buffer (lysis buffer supplemented with 1 M KCl) and washed again with 3 column volumes of lysis buffer. The His_6_-Sumo-tagged protein was then step-eluted with elution buffer (cell lysis buffer supplemented with 0.5 M imidazole) and added SENP2 protease to a concentration of 50 μg/mL (produced in-house by the Protein Technologies core facility at the CRG) to remove the tag. The digestion proceeded overnight at 4°C while simultaneously dialyzing the mixture against 4 L of cell lysis buffer (no imidazole) previously refrigerated using a 3.5 kDa cut-off Spectra/Por membrane from Thermo Fisher. The sample was then loaded again on a 5 mL His-Trap Fast Flow column and the flow-through was concentrated down to 0.5 mL using a 15 mL Amicon centrifugal filter with a 3.5 kDa cutoff (Merck). The sample was then loaded on a Superdex 75 Increase 10/300 GL column mounted in the same chromatography system and equilibrated in NMR buffer (20 mM sodium phosphate, 100 mM NaCl, pH 7.0). Fraction purity was assessed by SDS-PAGE using precast NuPAGE Bis-Tris gels from ThermoFisher, and fractions with >95% purity were pooled. Samples were quantified based on their calculated extinction coefficients and absorbance at 280 nm measured in a NanoDrop One instrument (Thermo Fisher), aliquoted, flash frozen in liquid nitrogen and stored at −80°C.

### NMR and CD spectroscopy

Recombinant protein sample aliquots were thawed on ice on data acquisition day, added 10% D_2_O (Eurisotop) for deuterium channel lock and 10 μM DSS (3-(Trimethylsilyl)-1-propanesulfonic acid sodium salt, Merck) for internal chemical shift referencing, and loaded in 3 mm NMR tubes (Norell, Morganton, NC, US). NMR spectra were acquired at 303 K (30°C, same temperature as variant selection assays) on a Bruker Avance III 600 MHz spectrometer equipped with a TCI cryoprobe and using TopSpin 4.0.8 for data acquisition and processing (Bruker, Billerica, MA, US). 2D ^1^H-^15^N heteronuclear correlation FAST-HSQC ^82^ spectra were acquired with 8 accumulations, a 1 s inter-scan delay and 2048 points in the direct ^1^H dimension and 256 in the indirect ^15^N dimension. Spectra were analyzed using CcpNmr Analysis version 2.5.2 ^83^.

Circular dichroism spectra were acquired on ^15^N-labeled recombinant protein samples diluted in NMR buffer and loaded in a 1 mm optical path quartz cuvette (Hellma, Müllheim, Germany) using a Jasco 815 UV circular dichroism spectrophotometer. 20-accumulation spectra were acquired in the 190-260 nm wavelength range with a 0.2 nm data interval and a scanning speed of 50 nm/min. For each sample, first a spectrum was acquired at 303 K. Then, samples were exposed to thermal denaturation via a temperature ramp in the 303-368 K range at a 2.5 degree per minute rate and ellipticity sampling at 222 nm every 0.5 degrees. Finally, a 3-accumulation spectrum with all other parameters set as detailed above was acquired at 368 K. Blank measurements were acquired at each experimental condition and subtracted from the sample data. Univariate splines were fit to the CD spectra and sigmoids were fit to the thermal denaturation data using the Python Scipy package. Melting temperature (T_m_) was obtained as the T value at the minimum of the first derivative of the sigmoid fit to the thermal melting data.

## Supplementary figures

**Fig S1.**
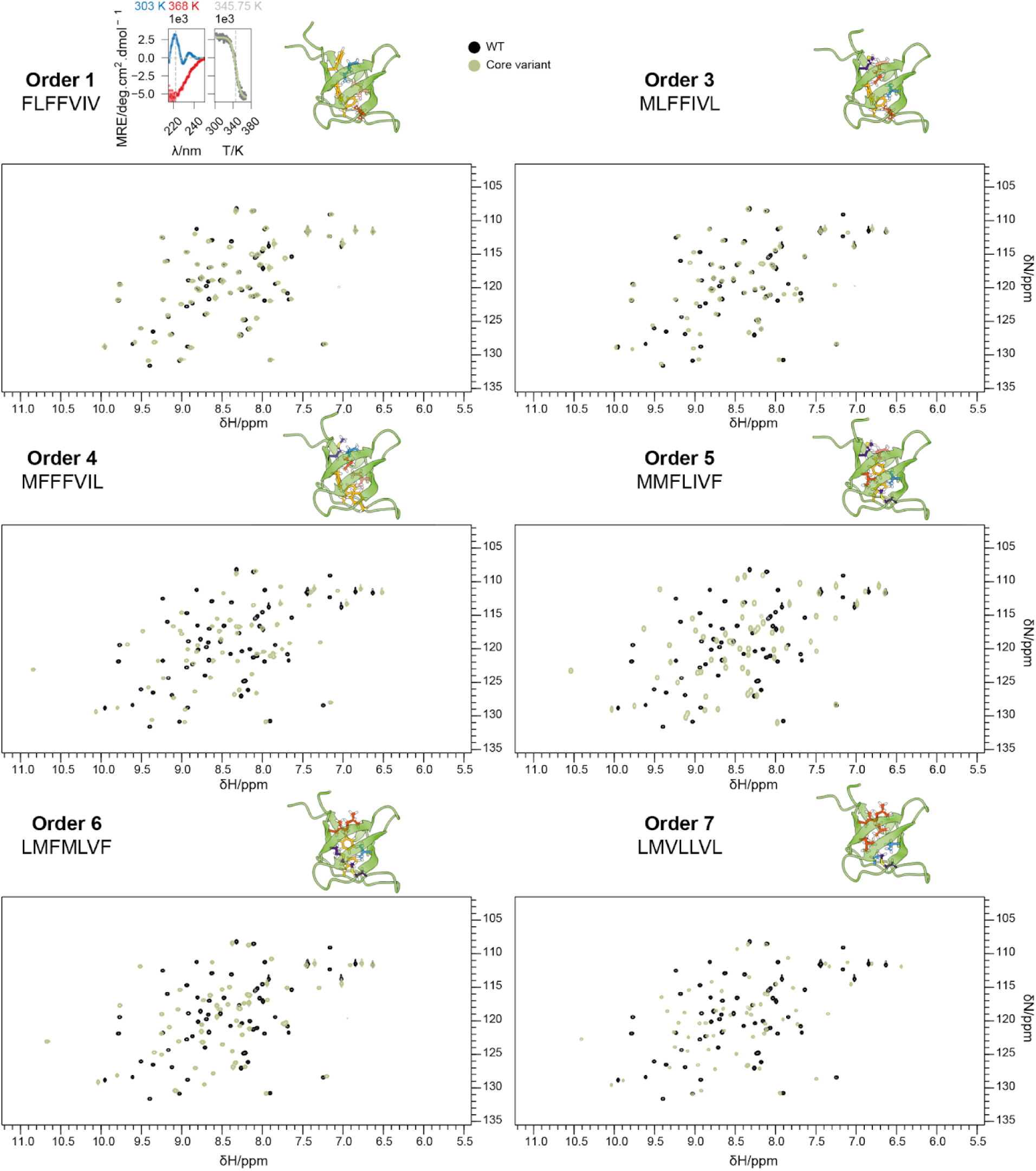
Comparison of the FYN-SH3 WT and core variants ^1^H-^15^N HSQC spectra. Spectra acquired on the high fitness, high order core variants (variant selection in Fig 1e.) are shown in green overlaid to the spectrum acquired on the WT (black). CD data is shown for a first order mutant as in Fig 2a.

**Fig S2.**
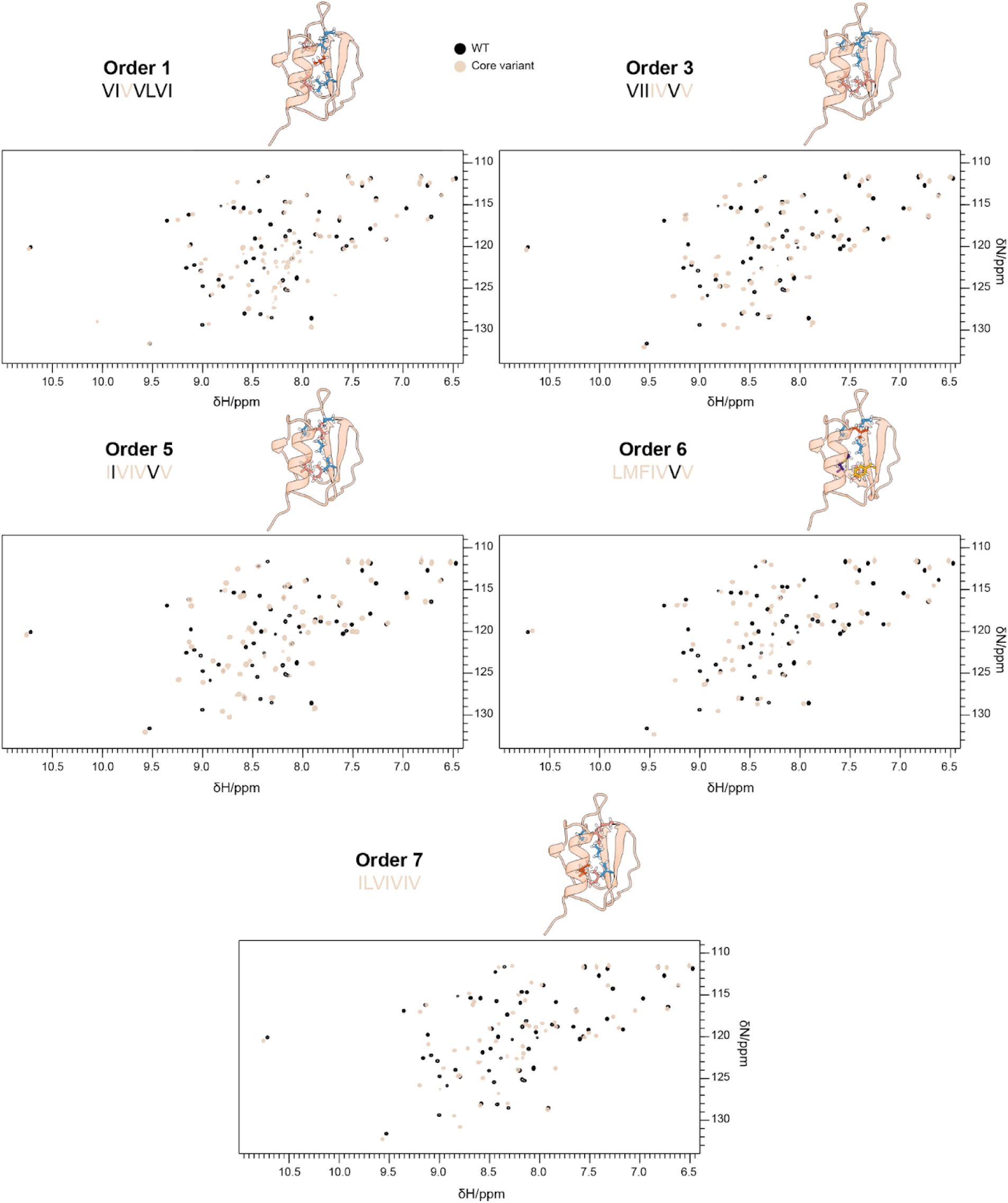
Comparison of the CI-2A WT and core variants ^1^H-^15^N HSQC spectra. Spectra acquired on the high fitness, high order core variants (variant selection in Fig 1e.) are shown in pink overlaid to the spectrum acquired on the WT (black).

**Fig S3.**
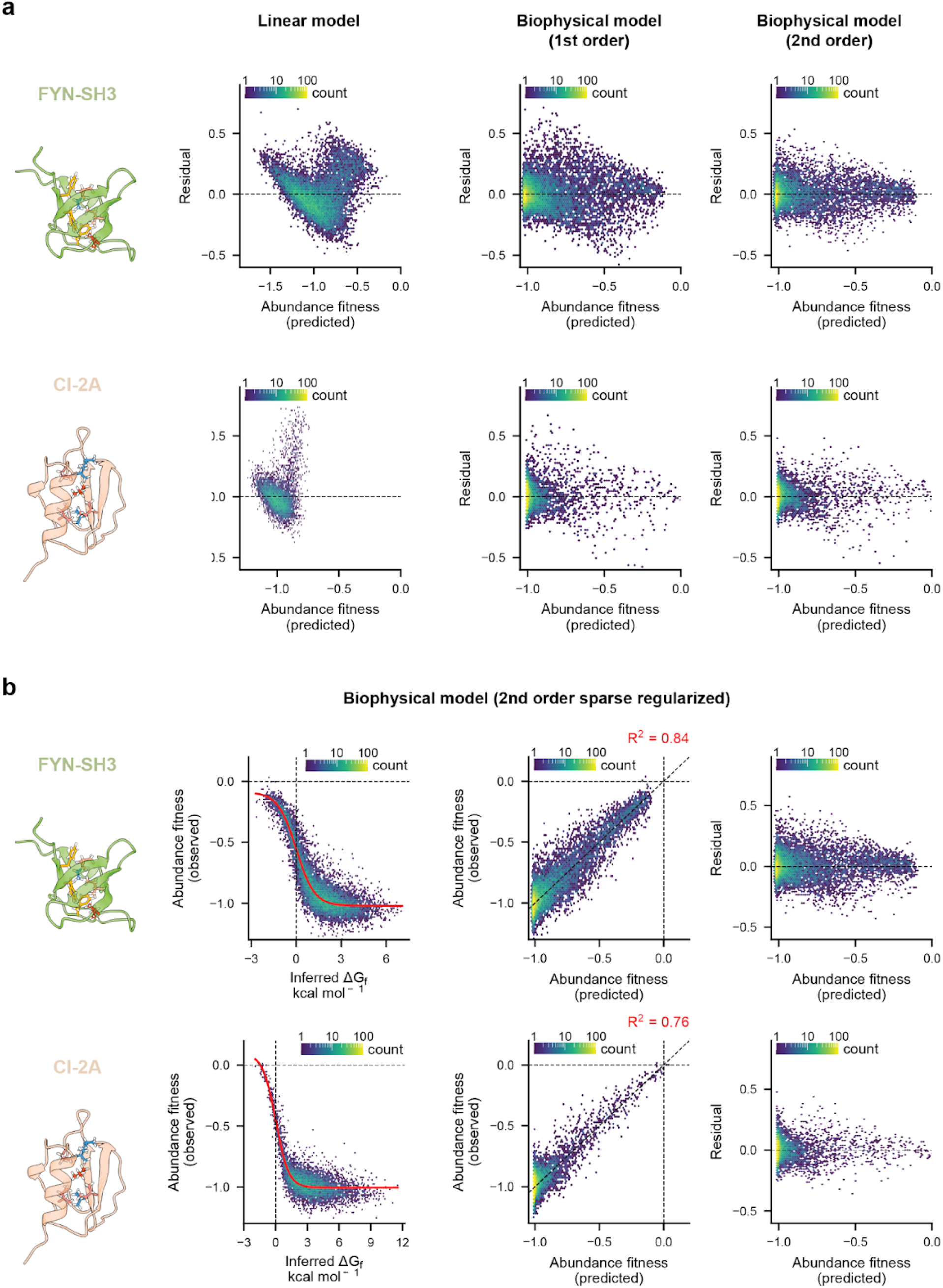
Supporting data on the models fit to the abundance fitness core mutagenesis datasets. **a.** Residuals plots for the linear and energy models shown in Fig 3. **b.** Sparse regularized second-order model fit on the FYN-SH3 and CI-2A core DTS mutagenesis datasets along with model performance and residuals.

**Fig S4.**
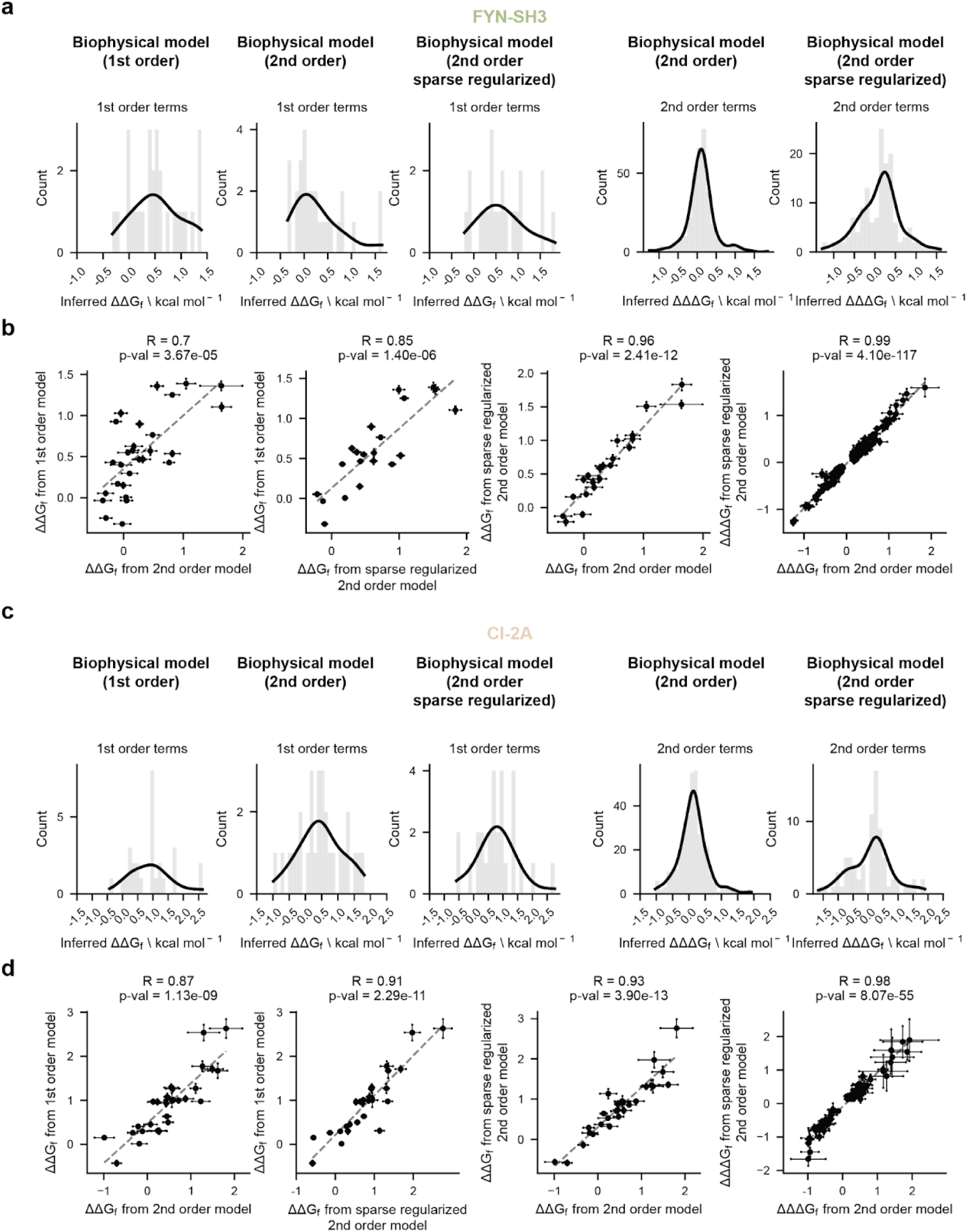
Comparison of energy terms inferred by all models fit to the abundance fitness core mutagenesis datasets. **a.** Histograms and density plots of the first (ΔΔG_f_) and second (ΔΔΔG_f_) order terms inferred by the models fit to the FYN-SH3 core DTS mutagenesis abundance fitness dataset. **b.** Relationship between the terms inferred by the different models for the FYN-SH3 core DTS mutagenesis abundance fitness dataset. **c.** As in **a.** for the CI-2A core DTS mutagenesis abundance fitness dataset. **d.** As in **b.** for CI-2A.

**Fig S5.**
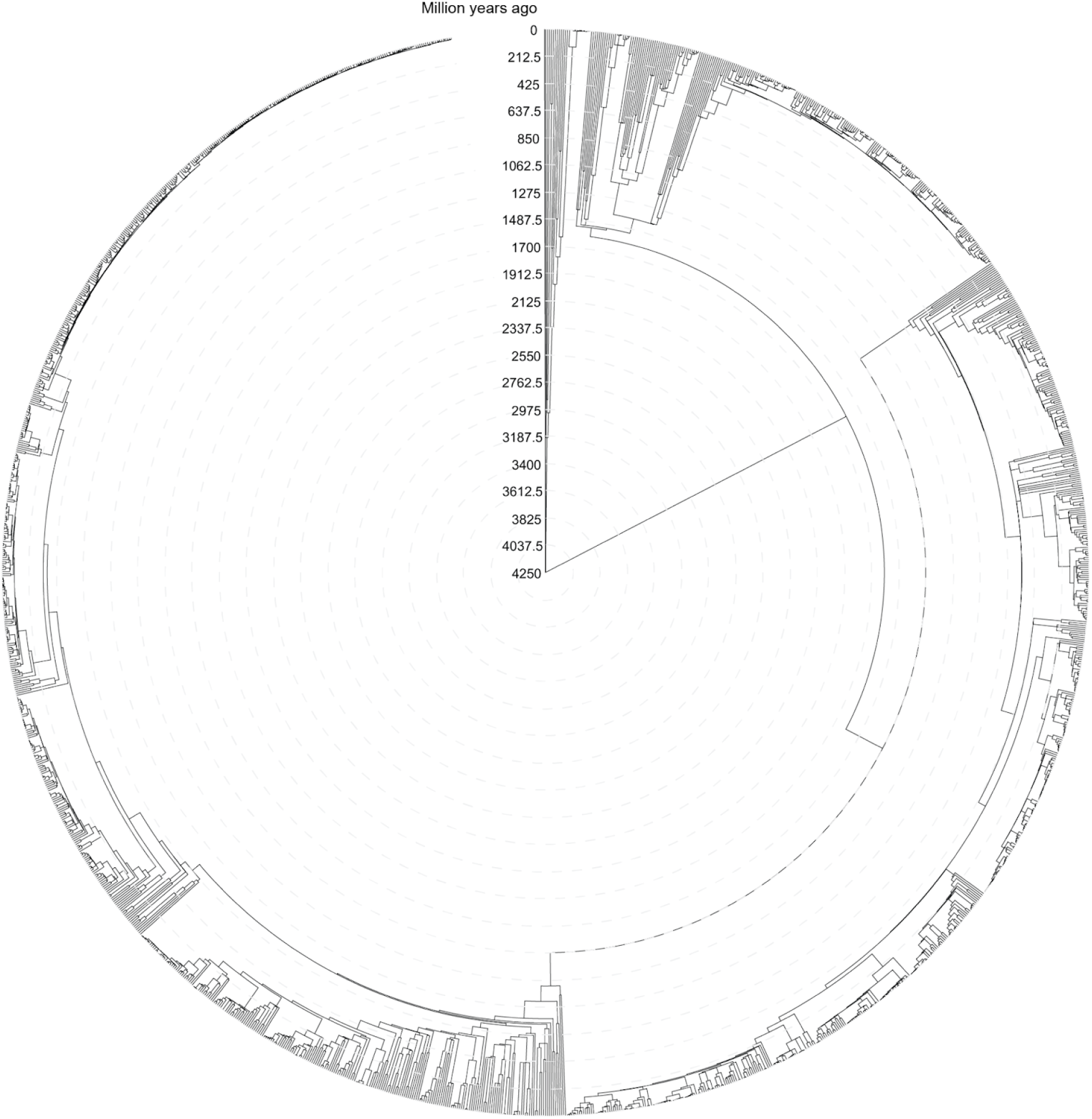
Phylogenetic tree of species with representation in the FYN-SH3 structural homologs set. The complete list of 2,545 species was uploaded to TimeTree 5 ^68^ (www.timetree.org), which mapped and provided time divergence estimates for 1,648 of them. The tree was visualized in iTOL v6 ^84^ (www.itol.embl.de). Gray dashed contours represent time divergence estimates in million years as indicated by the scale.

**Fig S6.**
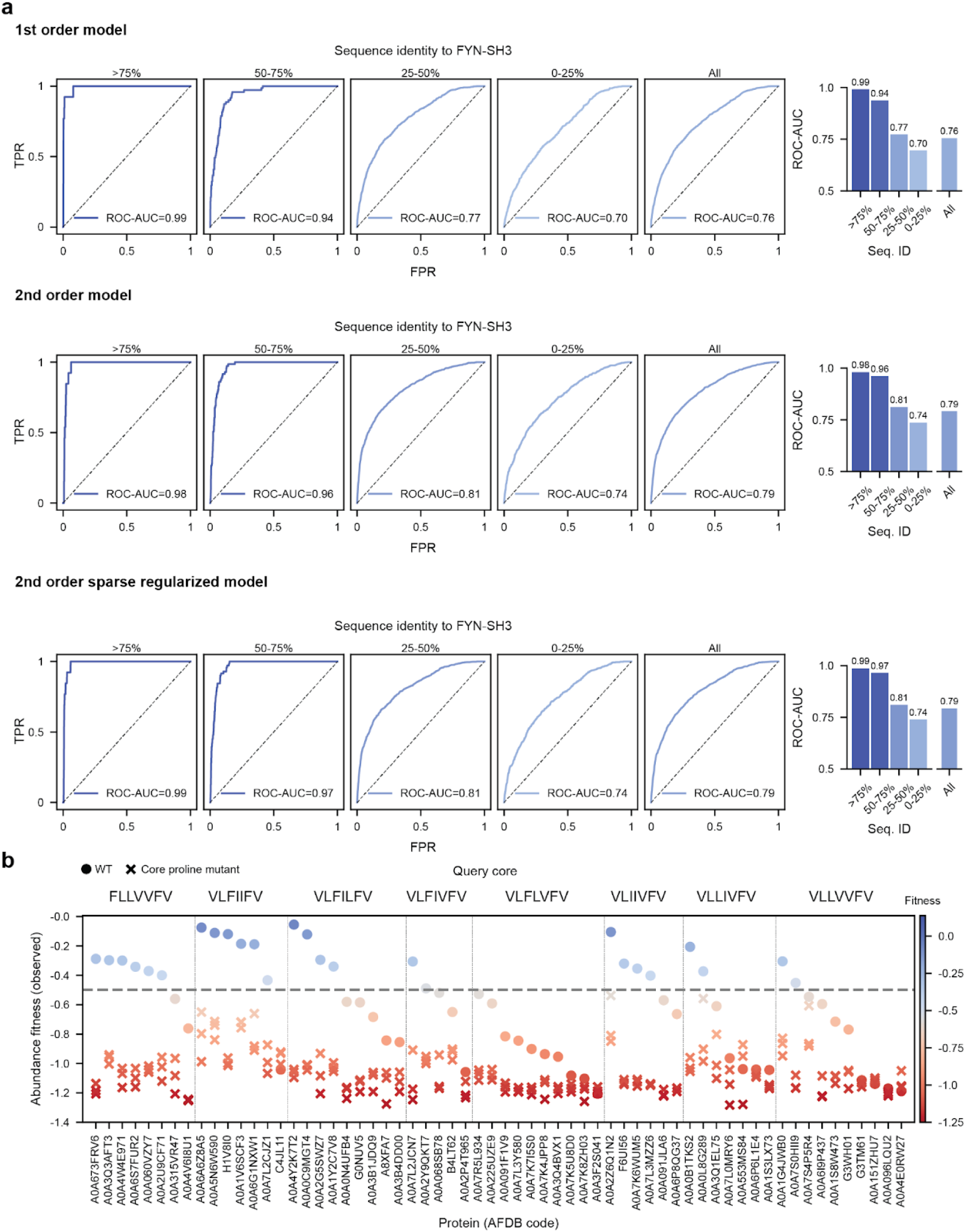
Supporting data on Earth Biogenome structural homolog classification and stability. **a.** Performance of all energy models fit to the FYN-SH3 core DTS mutagenesis abundance fitness dataset at classifying SH3 domains from the Earth Biogenome identified by structural homology. **b.** Abundance fitness of randomly picked structural homologs from the Earth Biogenome carrying hydrophobic cores that are detrimental when transplanted in FYN. WT sequences as dots and 3 proline core mutants per protein as crosses. Single proline mutations were randomly introduced in core positions.

**Fig S7.**
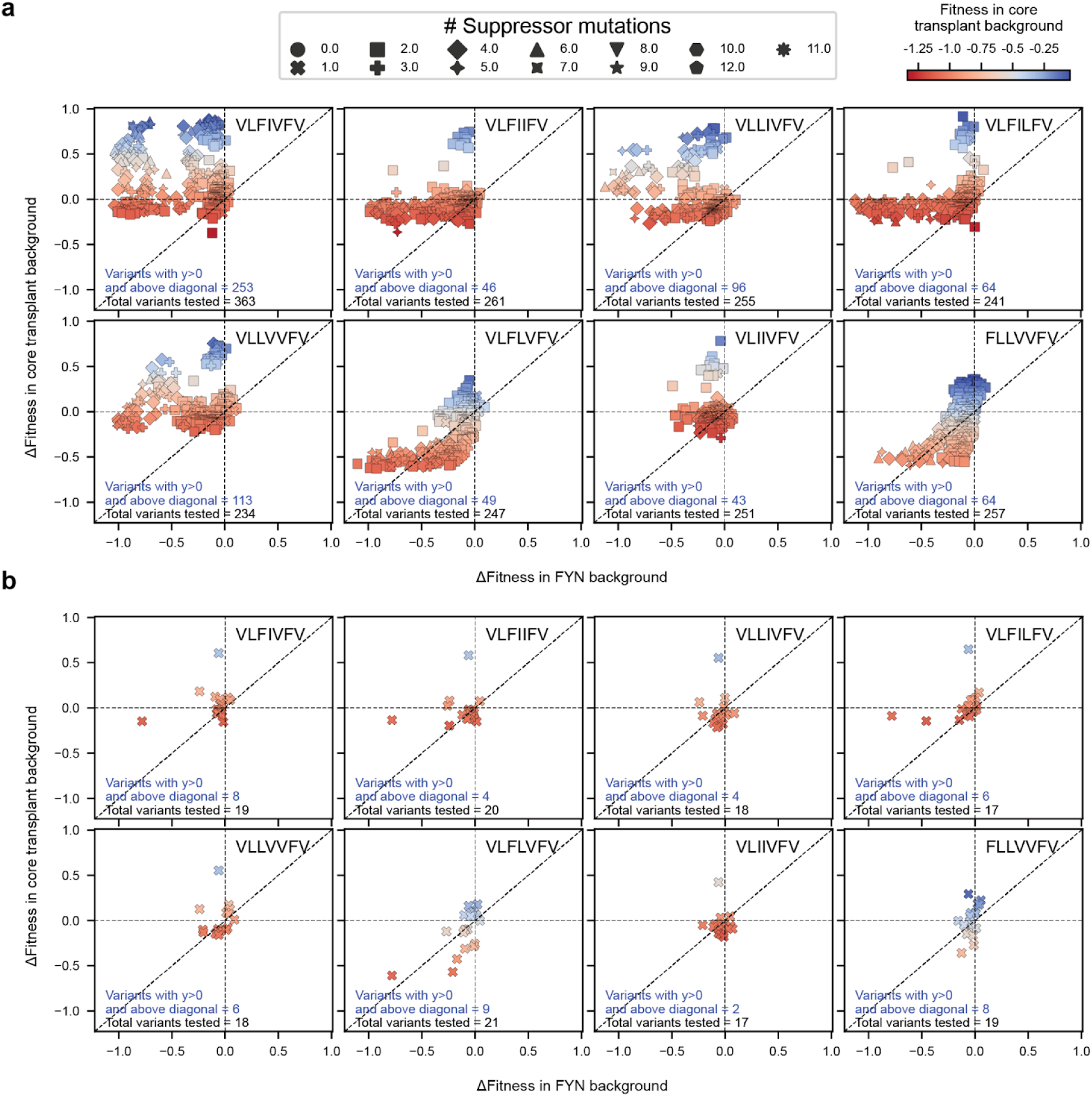
Experimentally testing candidate core transplantation failure suppressors. **a.** Difference in fitness for the variant with suppressor mutations of all orders in the WT *versus* the core transplant backgrounds. Variants above the horizontal and diagonal lines improve fitness in the core transplant background more than in the WT background. **b.** As in **a.** for variants with single suppressor mutations.

**Fig S8.**
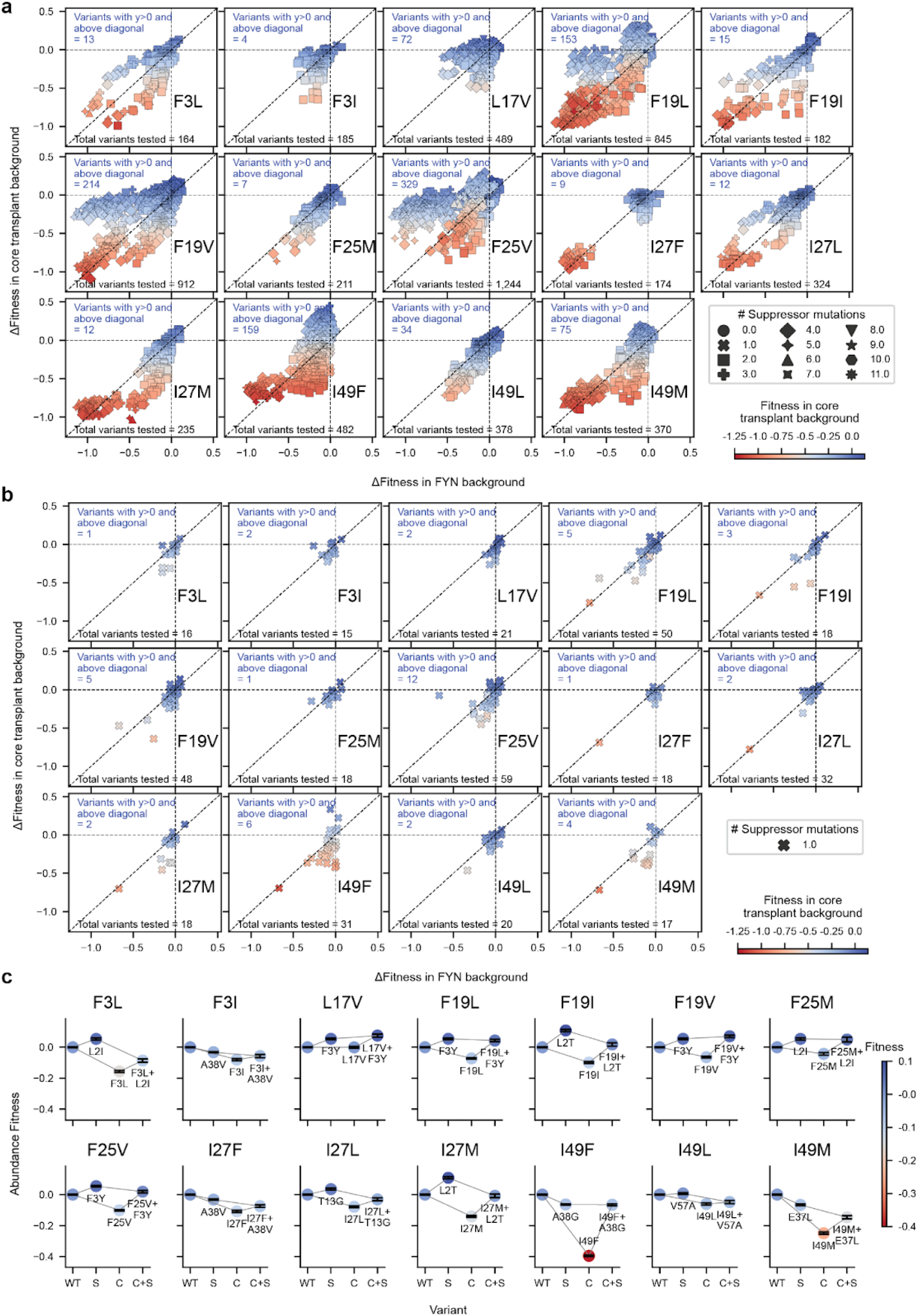
Detrimental FYN-SH3 single core mutation suppressors. **a.** Difference in fitness for the variant with suppressor mutations of all orders in the WT *versus* the single core mutant backgrounds. Variants above the horizontal and diagonal lines improve fitness in the single core mutant background more than in the WT background. **b.** As in **a.** for variants with single suppressor mutations. **c.** Best single suppressor mutation for each mildly detrimental single core mutation. *WT*, fitness of the wild-type. *S*, fitness of the suppressor mutant on the WT background. *C*, fitness of the single core mutant. *C+S*, fitness of the variant carrying both the core and suppressor mutations.

**Fig S9.**
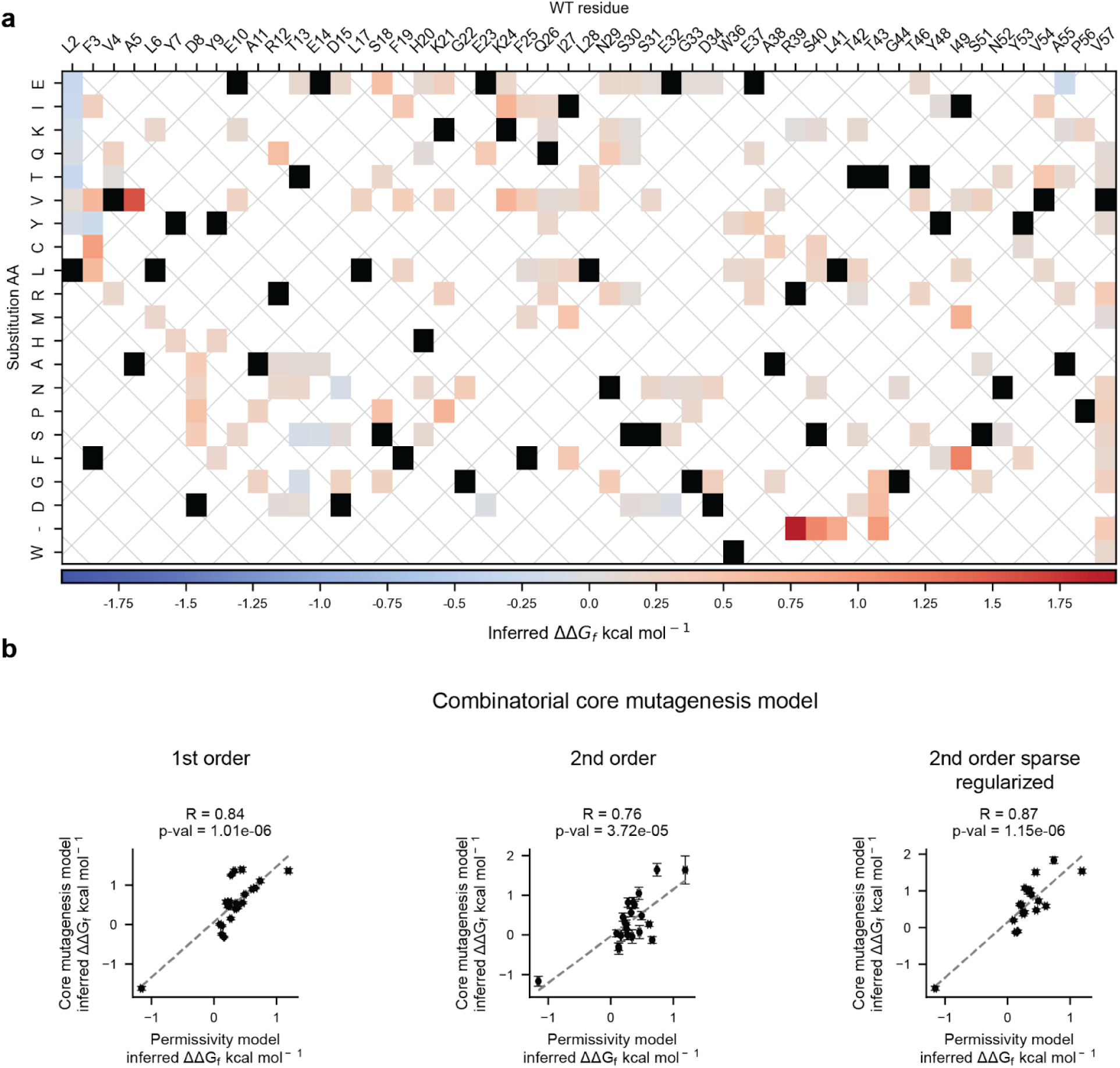
Folding free energy changes (*ΔΔG_f_*) inferred by the second-order sparse regularized model fit to the FYN suppression dataset. **a.** Heatmap of the sparse (*ΔΔG_f_*) terms inferred by the model. Black squares denote the WT amino acid at each position. **b.** Relationship of the terms in **a.** with equivalent terms inferred by the energy models fit on the FYN-SH3 core DTS mutagenesis abundance fitness dataset.

**Fig S10.**
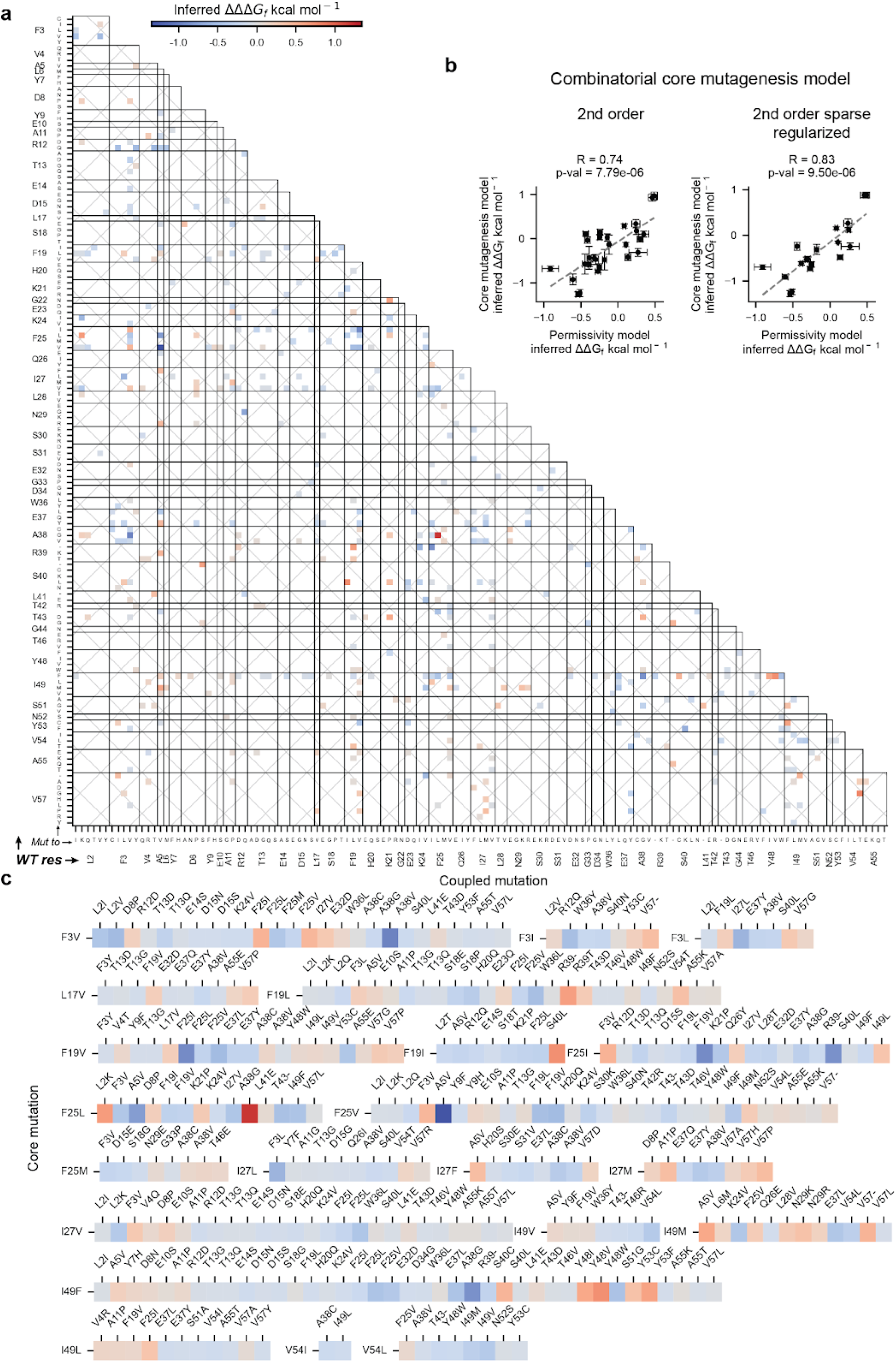
Pairwise energy couplings (*ΔΔΔG_f_*) in the second-order sparse regularized model fit to the FYN suppressor dataset. **a.** Heatmap of the sparse (*ΔΔΔG_f_*) pairwise energy coupling terms inferred by the model. **b.** Relationship of the terms in **a.** with equivalent terms inferred by the second-order energy models fit on the FYN-SH3 core DTS mutagenesis abundance fitness dataset. **c.** Pairwise energy couplings (*ΔΔΔG_f_*) involving core residues.

**Fig S11.**
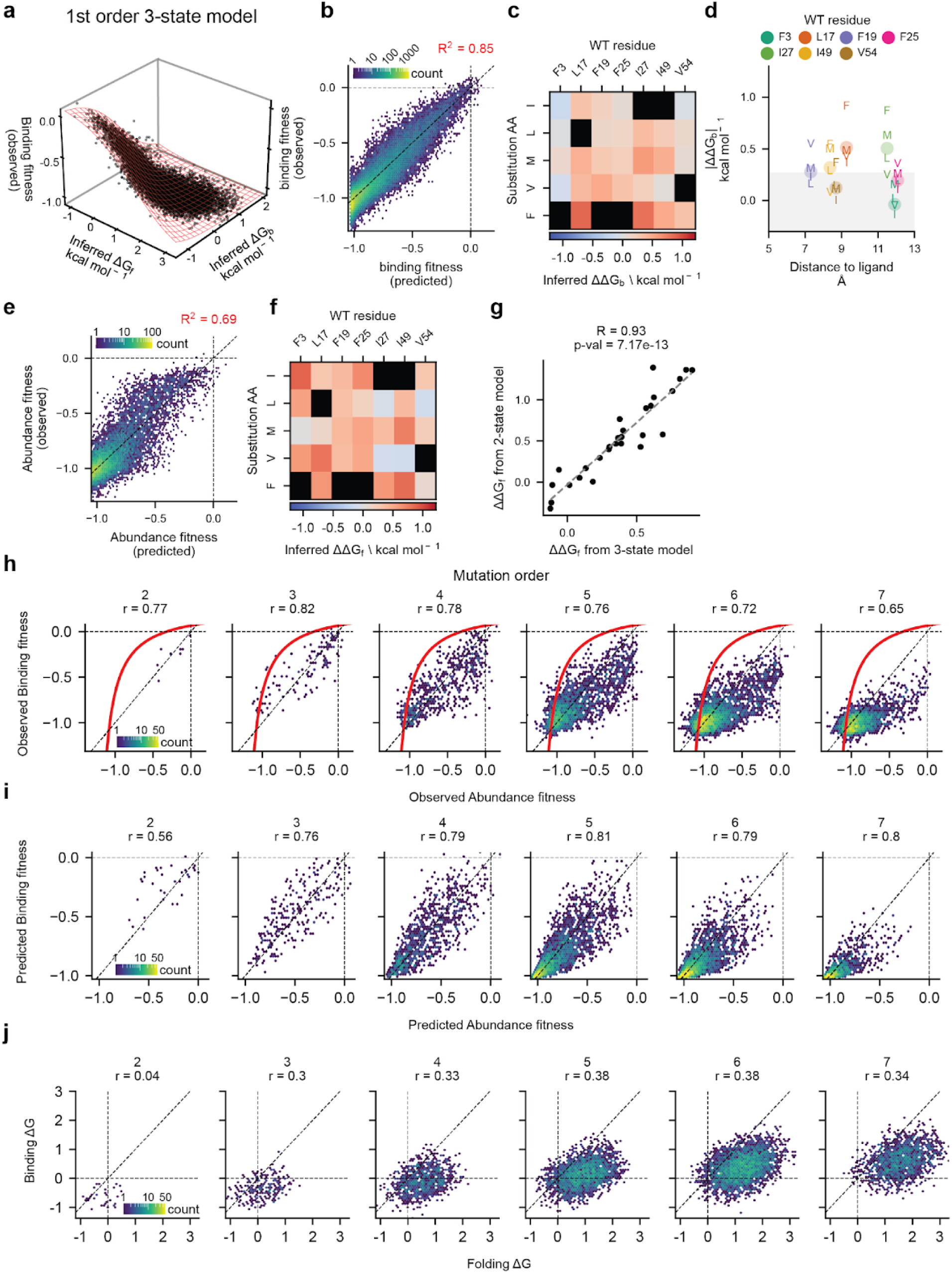
Details of the first order 3-state model of FYN-SH3 folding and binding to PRD1-super. **a.** Relationship between the observed binding fitness values and the inferred free energies of folding and binding (data as black dots, sigmoidal fit shown as a red mesh). **b.** Genetic prediction performance of the first order three-state model on binding fitness. **c.** Genetic background-averaged binding free energy changes (ΔΔG_b_) associated with FYN-SH3 core point mutations inferred by the first order three-state model. **d.** Magnitude of individual (characters) and position averaged (dots) single mutational binding free energy effects (|ΔΔG_b_|) versus minimum side chain heavy atom distance to ligand. **e.** Genetic prediction performance of the first order three-state model on abundance fitness. **f.** Genetic background-averaged folding free energy changes (ΔΔG_f_) associated with FYN-SH3 core point mutations inferred by the first order three-state model. **g.** Relationship between folding free energy changes (ΔΔG_f_) inferred by the two and three-state first order models fit on abundance fitness data. **h.** Relationship between abundance and binding fitness experimentally measured on FYN-SH3 core variants. Red lines indicate first order model-derived binding fitness expectancy based solely on abundance fitness (assuming no changes in binding free energy ΔΔG_b_). **i.** Relationship between binding and abundance fitness as predicted by the first order model for variants in the training set. **j.** Relationship between binding and folding free energies (ΔG_b_ and ΔG_f_) inferred by the first order model for variants in the training set as inferred by the model.

**Fig S12.**
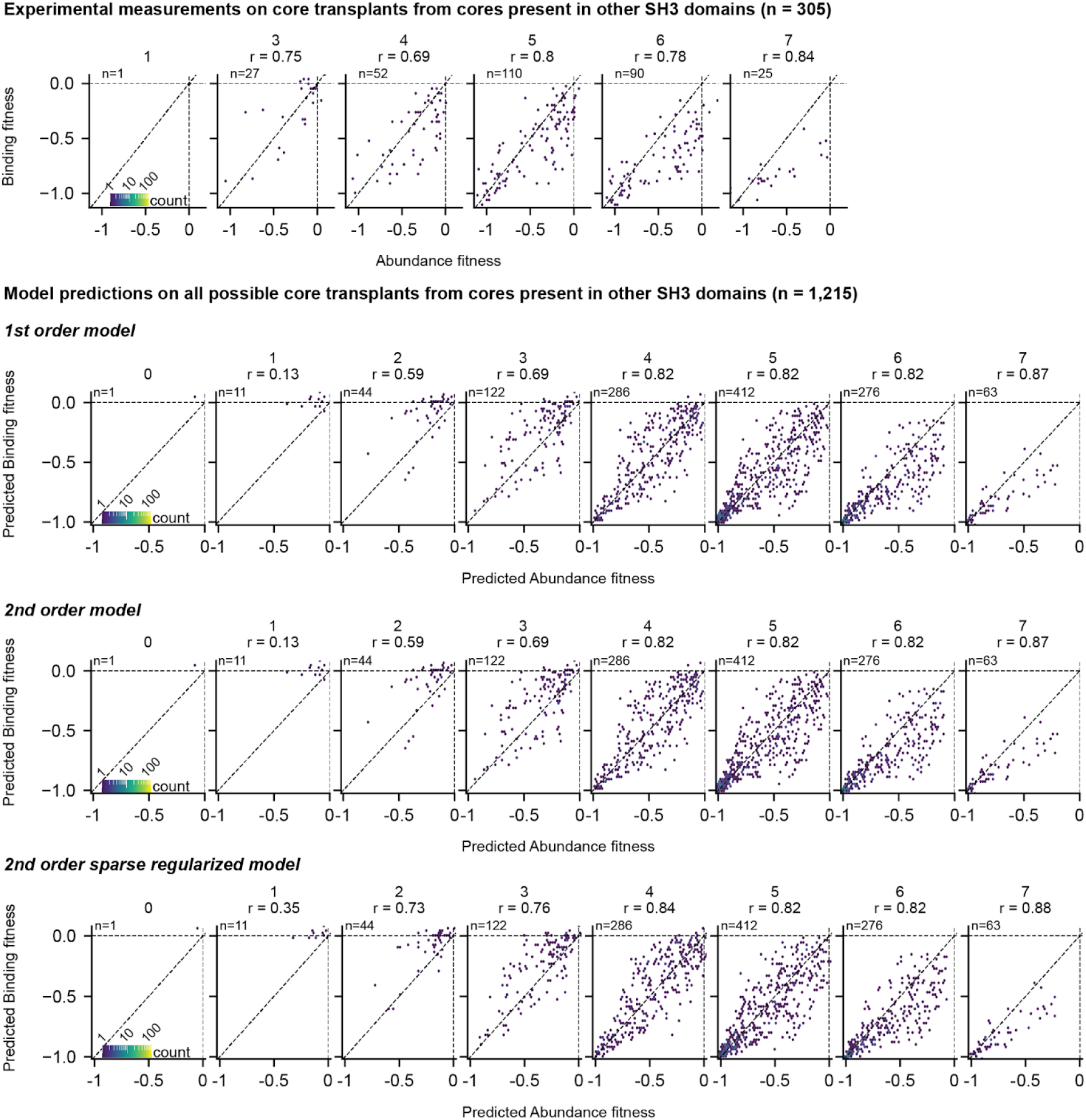
Relationship between abundance and binding fitness for FYN-SH3 variants carrying the cores of structural homologs in the Earth Biogenome. Top panel shows experimental data observed in the abundance and binding selections of the FYN-SH3 core DTS mutagenesis library. Other panels show predictions from the three-state models simultaneously fit on both datasets.

**Fig S13.**
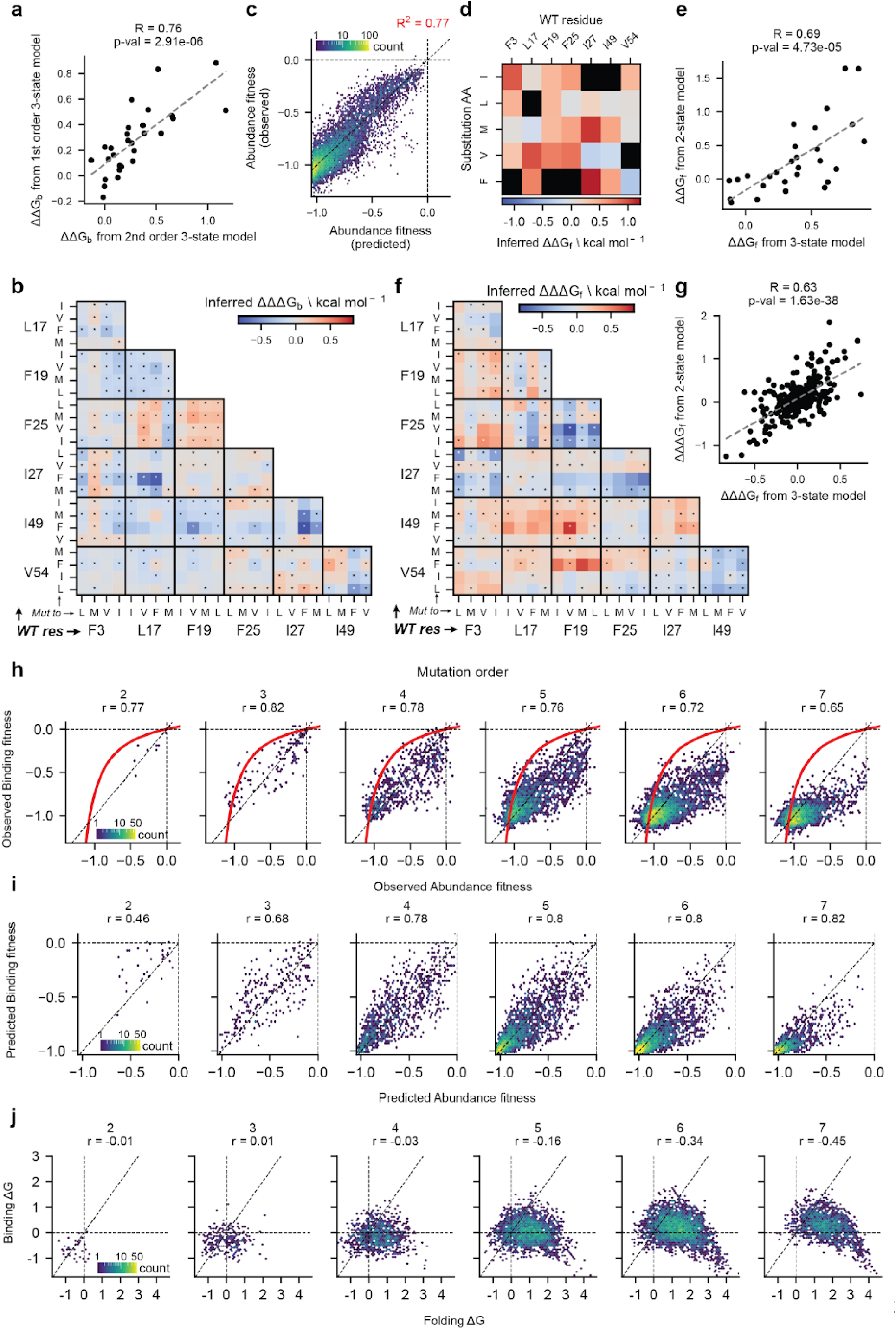
Details of the second-order 3-state model of FYN-SH3 folding and binding to PRD1-super. **a.** Relationship between binding free energy changes (ΔΔG_b_) inferred by the first and second-order three-state models fit simultaneously on binding and abundance fitness data. **b.** Pairwise binding energy couplings (ΔΔΔG_b_) inferred by the model. **c.** Genetic prediction performance of the second-order three-state model on abundance fitness. **d.** Genetic background-averaged folding free energy changes (ΔΔG_f_) associated with FYN-SH3 core point mutations inferred by the second-order three-state model. **e.** Relationship between folding free energy changes (ΔΔG_f_) inferred by the two and three-state second-order models fit on abundance fitness data. **f.** Pairwise folding energy couplings (ΔΔΔG_f_) inferred by the model. **g.** Relationship between pairwise folding energy couplings (ΔΔΔG_f_) inferred by the two and three-state second-order models fit on abundance fitness data. **h.** Relationship between abundance and binding fitness experimentally measured on FYN-SH3 core variants. Red lines indicate second-order model-derived binding fitness expectancy based solely on abundance fitness (assuming no changes in binding free energy ΔΔG_b_). **i.** Relationship between binding and abundance fitness as predicted by the second-order model for variants in the training set. **j.** Relationship between binding and folding free energies (ΔG_b_ and ΔG_f_) inferred by the second-order model for variants in the training set as inferred by the model.

**Fig S14.**
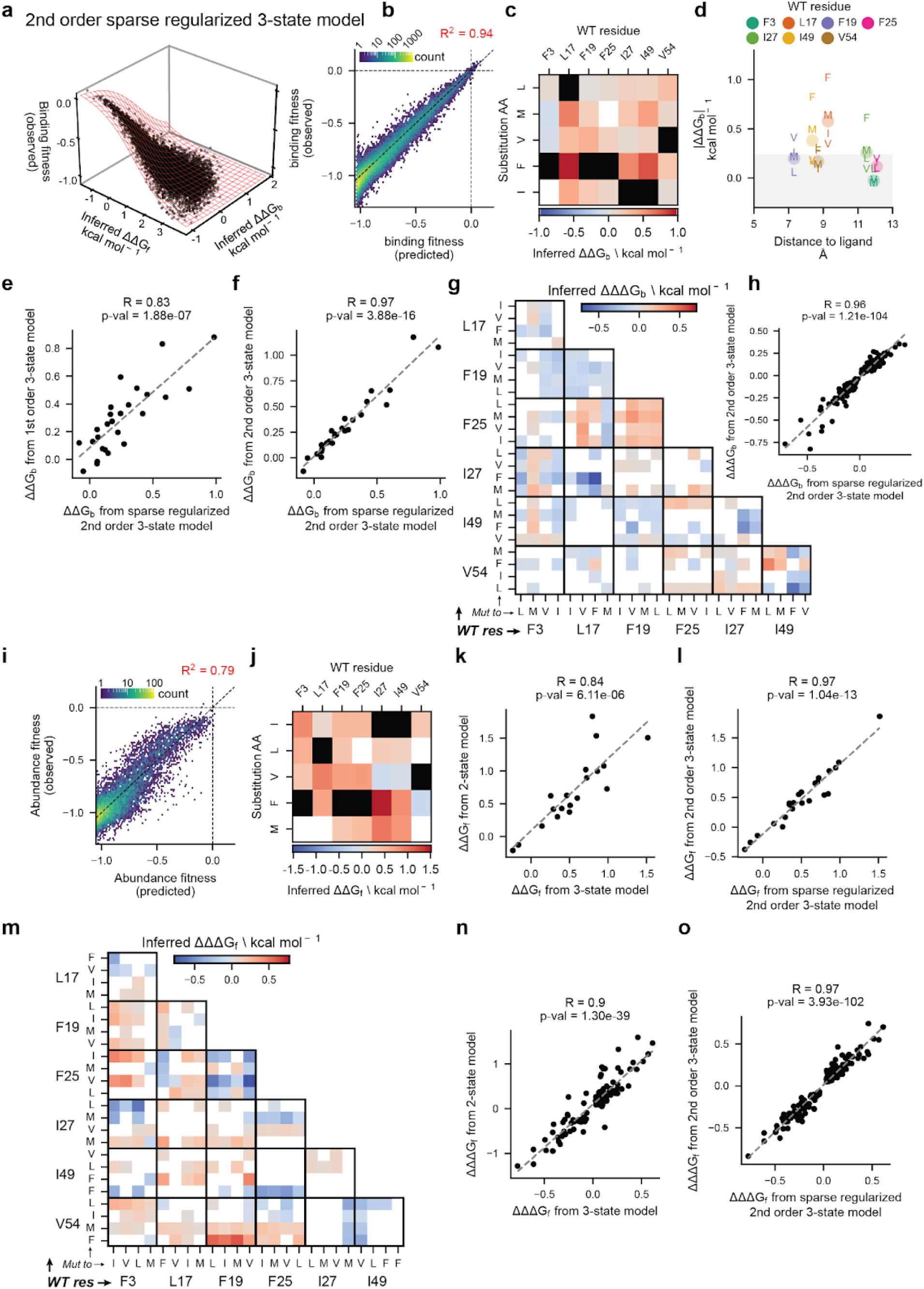
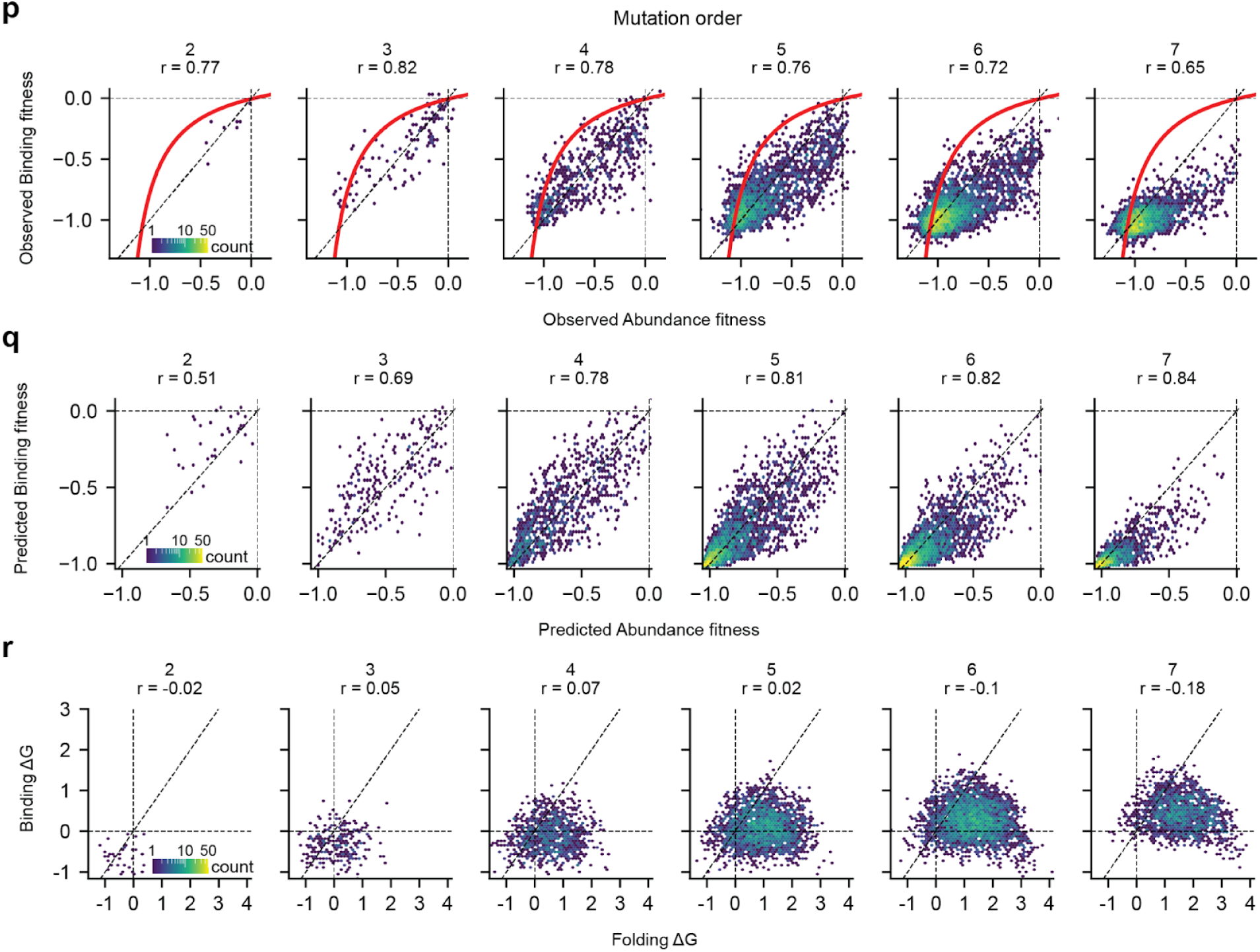
Details of the second-order sparse regularized 3-state model of FYN-SH3 folding and binding to PRD1-super. **a.** Relationship between the observed binding fitness values and the inferred free energies of folding and binding (data as black dots, sigmoidal fit shown as a red mesh). **b.** Genetic prediction performance of the second-order sparse regularized three-state model on binding fitness. **c.** Genetic background-averaged binding free energy changes (ΔΔG_b_) associated with FYN-SH3 core point mutations inferred by the second-order sparse regularized three-state model. **d.** Magnitude of individual (characters) and position averaged (dots) single mutational binding free energy effects (|ΔΔG_b_|) versus minimum side chain heavy atom distance to ligand. **e.** Relationship between the binding free energy changes (ΔΔG_b_) inferred by the first order and the second-order sparse regularized 3-state models. **f.** Relationship between the binding free energy changes (ΔΔG_b_) inferred by the full second-order and the second-order sparse regularized 3-state models. **g.** Pairwise binding energy couplings (ΔΔΔG_b_) inferred by the second-order sparse regularized 3-state model. **h.** Relationship between the pairwise binding energy couplings (ΔΔΔG_b_) inferred by the full second-order and the second-order sparse regularized 3-state models. **i.** Genetic prediction performance of the second-order sparse regularized three-state model on abundance fitness. **j.** Genetic background-averaged folding free energy changes (ΔΔG_f_) associated with FYN-SH3 core point mutations inferred by the second-order sparse regularized three-state model. **k.** Relationship between folding free energy changes (ΔΔG_f_) inferred by the two and three-state second-order sparse regularized models fit on abundance fitness data. **l.** Relationship between folding free energy changes (ΔΔG_f_) inferred by the full second-order and the second-order sparse regularized 3-state models fit on abundance fitness data. **m.** Pairwise folding energy couplings (ΔΔΔG_f_) inferred by the second-order sparse regularized 3-state model. **n.** Relationship between the pairwise folding energy couplings (ΔΔΔG_f_) inferred by the two and three-state second-order sparse regularized models fit on abundance fitness data. **o.** Relationship between pairwise folding energy couplings (ΔΔΔG_f_) inferred by the full second-order and the second-order sparse regularized 3-state models fit on abundance fitness data. **p.** Relationship between abundance and binding fitness experimentally measured on FYN-SH3 core variants. Red lines indicate second-order sparse regularized model-derived binding fitness expectancy based solely on abundance fitness (assuming no changes in binding free energy ΔΔG_b_). **i.** Relationship between binding and abundance fitness as predicted by the second-order sparse regularized model for variants in the training set. **j.** Relationship between binding and folding free energies (ΔG_b_ and ΔG_f_) inferred by the second-order sparse regularized model for variants in the training set as inferred by the model.

## Supplementary tables

**Table S1.**
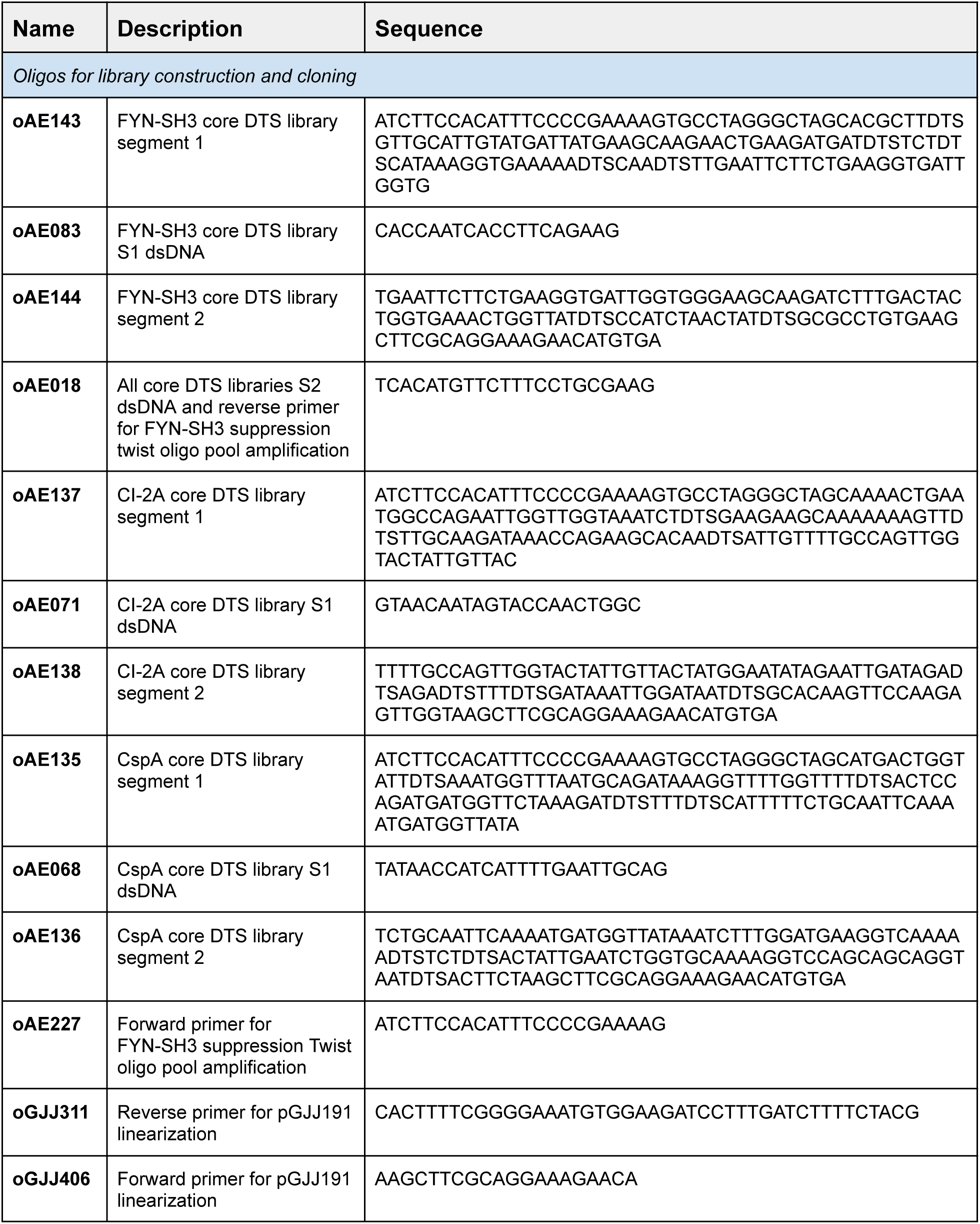
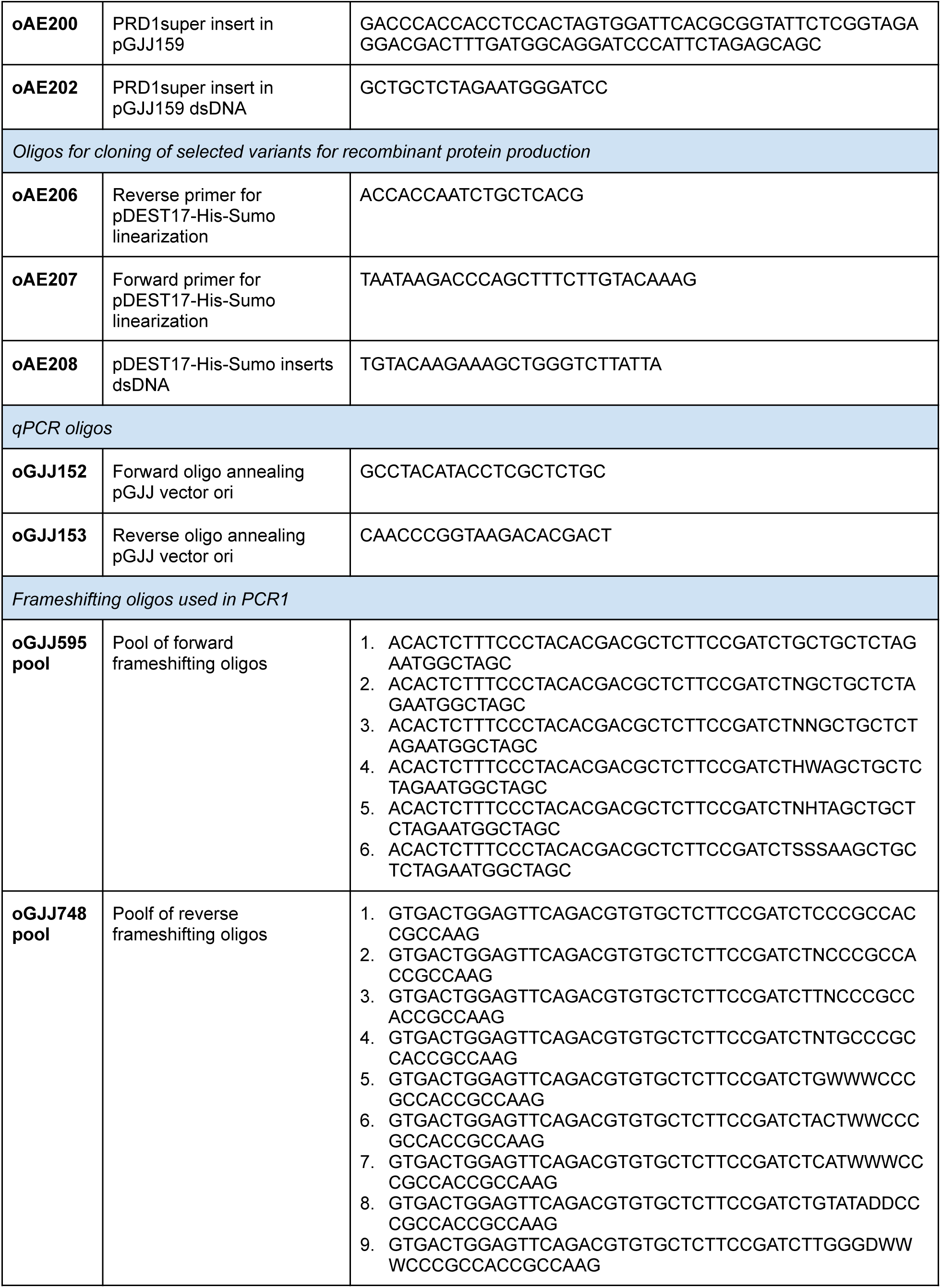
Oligonucleotides used in this study.

**Table S2.** Plasmids used in this study, available upon request, Material Transfer Agreement required.

| Name | Description | Benchling link |
| --- | --- | --- |
| pGJJ191 | Mutagenesis vector | https://benchling.com/s/seq-OajY23vrF4wB8WLA6xk8?m=slm-9rBpAQk5BglzfLp2qp3s |
| pGJJ159 | bindingPCA vector | https://benchling.com/s/seq-m88f9UhGMuEpzavHYzSk?m=slm-lDrFdggHI3KbgYVDNsiJ |
| pGJJ162 | abundancePCA vector | https://benchling.com/s/seq-nWq1p6ZFosa5PmRRTe1Q?m=slm-ajZ8Fr28gTx3UTKRvwCT |
| pDEST17-His-Sumo | Recombinant protein production in *E. coli* BL21(DE3) | https://benchling.com/s/seq-ZGBQn2M18iqj6y6MUmnf?m=slm-OOosjnzNZQBXYkdBEbwk |

**Table S3.**
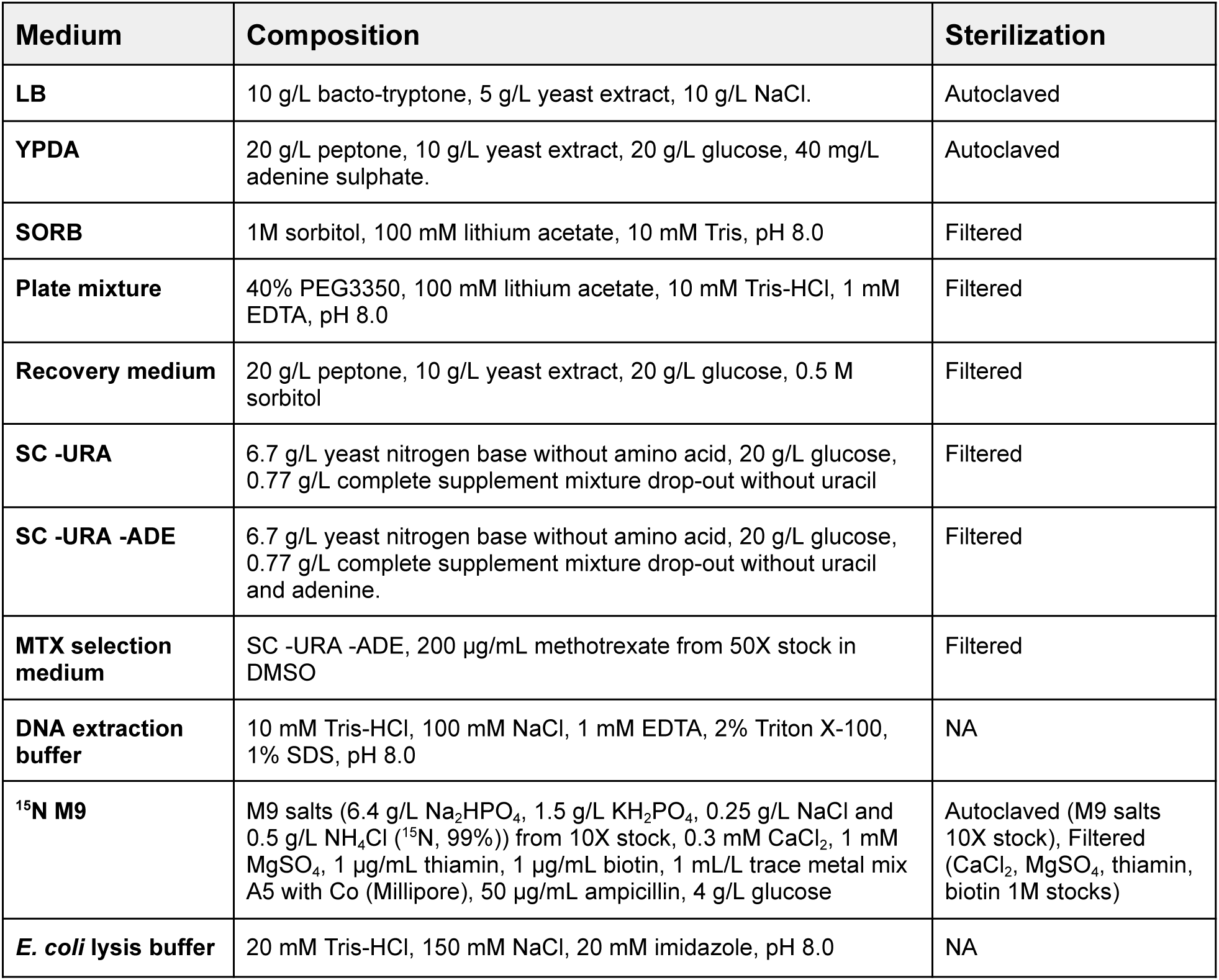
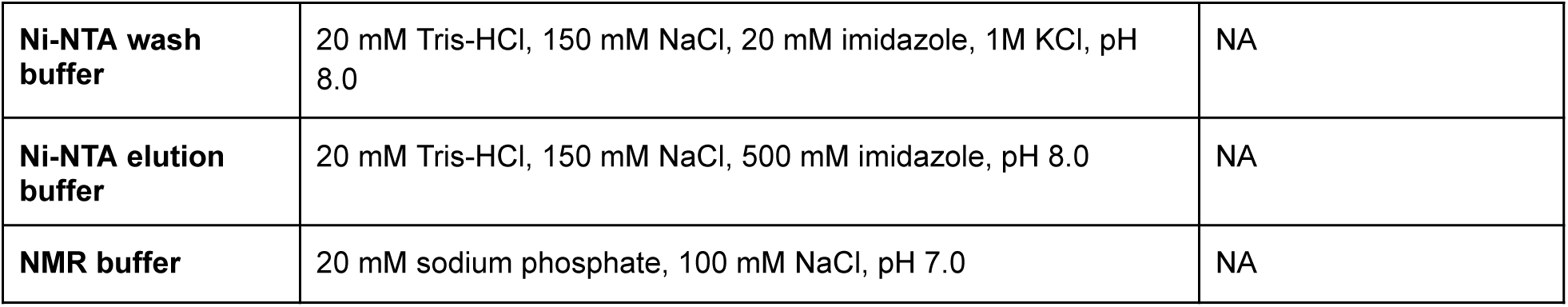
Non-commercial media and buffers used in this study. Autoclaving was performed at 120°C for 20 min. Filter sterilization was achieved using 0.2 mm Nylon membranes (Thermo Fisher).

